# A Natural qE- and qZ-Deficient Alga Retains Functional Plant-Like and Chlorophycean Violaxanthin De-Epoxidases

**DOI:** 10.64898/2026.09.26.754548

**Authors:** Karina Cunningham, Setsuko Wakao, Johanna Hall, Crysten E. Blaby-Haas, Rituparna Ivatury, Lucy Gleeson, Derrick Chuang, Jane J. Kim, Jeffrey L. Moseley, Graham R. Fleming, Krishna K. Niyogi

**Author notes:** **Corresponding author(s): Setsuko Wakao**, Molecular Biophysics and Integrated Bioimaging Division, Lawrence Berkeley National Laboratory, Berkeley, CA, USA.; Krishna K. Niyogi, Howard Hughes Medical Institute, University of California, Berkeley, CA, USA. The author responsible for distribution of materials integral to the findings presented in this article in accordance with the policy described in the Instructions for Authors (https://academic.oup.com/plphys/pages/General-Instructions) is Krishna K. Niyogi.

## Abstract

Photosynthetic organisms protect themselves from excess light through non-photochemical quenching (NPQ), a process that safely dissipates excess light energy as heat. Current understanding of NPQ is largely drawn from a few model systems, leaving it unclear how universal NPQ mechanisms are across the green lineage. We characterize the slow, sustained NPQ of the green alga, *Auxenochlorella protothecoides* x *symbiontica* (UTEX 250-A). The *Auxenochlorella* genome lacks Light-Harvesting Complex Stress Related (LHCSR), one of the two proteins required for the fast energy-dependent quenching (qE) component of NPQ. Mutants lacking the Photosystem II Subunit S (PSBS)-like gene (*psbsl1*) in *Auxenochlorella* had no effect on NPQ, indicating this alga lacks canonical qE. We identified two candidate enzymes associated with zeaxanthin-dependent quenching (qZ), which has slower kinetics: a plant-type Violaxanthin De-Epoxidase (VDE) and an algal-type Chlorophycean Violaxanthin De-Epoxidase (CVDE). High light did not trigger the expected conversion of violaxanthin to zeaxanthin, a pigment change that is a hallmark of qZ. Mutants lacking VDE (*vde1*), CVDE (*cvde1*), or both enzymes (*vde1 cvde1*) still displayed slow, reversible NPQ. However, when the *Auxenochlorella* VDE1 and CVDE1 were expressed in a VDE-deficient mutant of *Nicotiana benthamiana,* both enzymes restored zeaxanthin production and NPQ, confirming they are catalytically functional de-epoxidases. Together, these results reveal an alga that retains two functional, evolutionarily distinct xanthophyll-cycle enzymes yet does not rely on typical xanthophyll pigment dynamics for photoprotection. This points to a biological function that is decoupled from de-epoxidase activity, highlighting unexplored diversity in green algal photoprotection.

## Introduction

Photosynthetic organisms experience fluctuating light conditions on timescales of seconds to seasons. When the rate of excitation of the photosystems exceeds the capacity for photochemistry, excess excitation can increase the probability of molecular oxygen being converted to singlet oxygen (^1^O_2_*). ^1^O_2_* is a highly reactive form of reactive oxygen species (ROS) that directly oxidizes cell membranes, proteins, and DNA, and is considered the main cause of PSII photodamage (Triantaphylidès et al., 2008). To minimize photodamage, photosynthetic organisms have evolved photoprotective pathways that can prevent production of ROS by quenching surplus excitation energy and releasing it as heat, collectively measured and referred to as non-photochemical quenching (NPQ) of chlorophyll (Chl) fluorescence (Müller et al., 2001; Ruban et al., 2012; Niyogi and Truong, 2013). Although rapid activation of NPQ under excess light is crucial in preventing photodamage, the dissipation of excitation energy by NPQ directly competes with photosynthetic efficiency. As a result, NPQ must be rapidly turned off upon transitions from excess light to limiting light (Kromdijk et al., 2016; Long et al., 2022) and is fine-tuned through several mechanisms differing in timescale and quenching capacity.

NPQ consists of several components, including energy-dependent quenching (qE) and zeaxanthin-dependent quenching (qZ). qE develops within seconds to minutes in response to thylakoid lumen acidification and is mediated by pH-responsive proteins in the light-harvesting complex (LHC) superfamily, including Photosystem II Subunit S (PSBS) in plants (Li et al., 2000) and Light-Harvesting Complex Stress Related (LHCSR) in green algae (Peers et al., 2009). In the model green alga, *Chlamydomonas reinhardtii* (hereafter *Chlamydomonas*), qE is largely dependent on LHCSR proteins, whereas PSBS is considered to have a transient role in establishing NPQ (Correa-Galvis et al., 2016). Gene expression of both *LHCSR* and *PSBS* is strongly upregulated in high-light stress (Mettler et al., 2014), in contrast to other *LHCs* that are downregulated in high-light stress (Teramoto et al., 2002; Durnford et al., 2003; Elrad and Grossman, 2004).

The qE and qZ components of NPQ involve the conversion of violaxanthin (Vio), antheraxanthin (Anthera) and zeaxanthin (Zea) cycle (VAZ cycle) carotenoids (Yamamoto and Kamite, 1972; Demmig-Adams et al., 2014) (Fig. 1a and b). These three xanthophylls are oxygenated carotenoids that differ in the number of 5,6-epoxide groups on their terminal rings. Vio has two epoxides, Anthera has one, and Zea has none. De-epoxidation of Vio is catalyzed by violaxanthin de-epoxidase (VDE) in high light and reversed by zeaxanthin epoxidase (ZEP) in the dark (Fig. 1a). The interconversion of these pigments is mediated on a timescale of minutes to tens of minutes, which correlates with increased activation of qE and induction of slowly reversible qZ (Nilkens et al., 2010). The qZ component differs from Zea-dependent qE, because qZ no longer requires a low lumen pH once Zea has accumulated.

**Figure 1.**
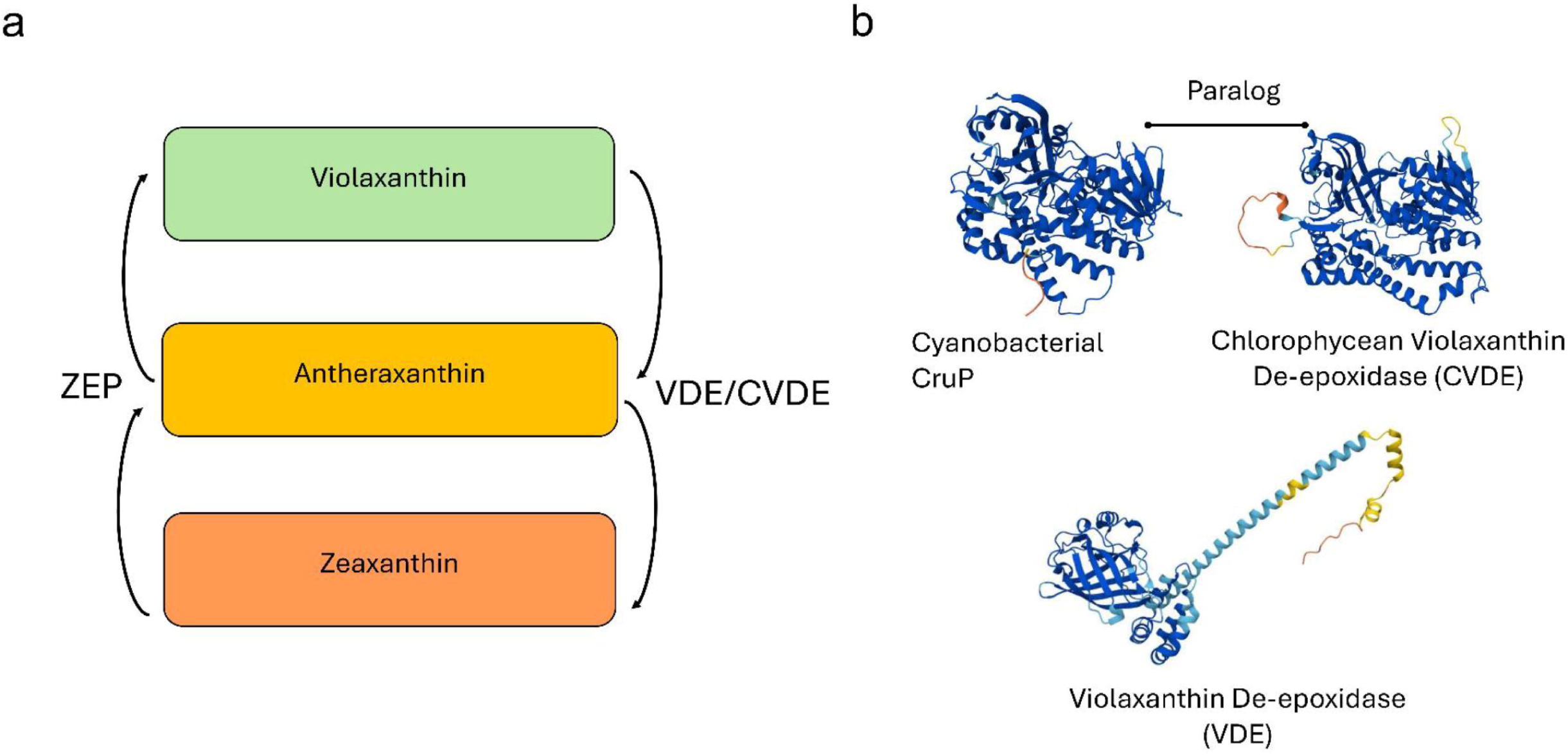
Graphical summary of VDE and CVDE in the biosynthetic pathway for xanthophyll cycle. **a)** Both CVDE and VDE enzymes convert violaxanthin to antheraxanthin and finally to zeaxanthin; the reverse conversion is mediated by zeaxanthin epoxidase (ZEP). **b)** Evolutionary relationship between cyanobacterial *Synechococcus sp. PCC 7002* CruP and chlorophycean violaxanthin de-epoxidase (CVDE) from *Chlamydomonas reinhardtii* and the *Arabidopsis thaliana* violaxanthin de-epoxidase (VDE).

Two evolutionarily and structurally distinct types of VDE are known. The plant-type VDE is a lipocalin-domain-containing enzyme localized to the thylakoid lumen (Bugos et al., 1998) (Fig. 1a). VDE requires acidic conditions and the reducing cofactor ascorbate (ascorbic acid) for activity (Hieber et al., 2002; Arnoux et al., 2009), and it is inhibited by the strong reducing agent dithiothreitol (DTT) (Yamamoto et al., 1972). In contrast, some green algae possess an analogous enzyme, referred to as chlorophycean violaxanthin de-epoxidase (CVDE), which is associated with the thylakoid membrane on the stromal side and is a paralog of the CruP protein (Li et al. 2016) (Fig. 1a). Although the exact function of CruP remains unknown, it is a homolog of CruA, a cyanobacterial lycopene β-cyclase (Bradbury et al., 2012) (Fig. 1a). CVDE is hypothesized to be activated under more alkaline conditions in the stroma but is not inhibited by DTT and its cofactor is unknown (Li et al., 2016).

All oxygenic phototrophs have evolved mechanisms to overcome the dilemma of safely dissipating excess light energy to protect the photosystems while maximizing photosynthetic efficiency for growth. In addition to qE and qZ, other components of NPQ with longer timescales have been identified, including qT that involves state transitions, qH that involves a sustained antenna quenching component, and qI that is associated with photoinhibition and damage to the D1 protein of photosystem II (Malnoë et al., 2018). However, different algal lineages and ecotypes have evolved to utilize unique combinations of photoprotective mechanisms. Elucidating and comparing these adaptations can reveal the evolutionary history of conserved and lineage-specific features. Additionally, understanding the diversity of photoprotection can give insight into design principles that can improve stress tolerance in crops and industrially relevant algae.

The eukaryotic unicellular green microalga, *Auxenochlorella* UTEX 250-A (hereafter *Auxenochlorella*) belongs to the trebouxiophyceae class (Craig et al., 2025). Trebouxiophyceae includes members such as *Picochlorum*, *Botryococcus*, and *Chlorella sp*, which have been used as feedstocks for biofuels (Anto et al., 2024; Ashokkumar et al., 2024). Some *Chlorella* species, such as *Chlorella ohadii*, are known for extreme high light and abiotic stress tolerance (Levin et al., 2021). *Auxenochlorella* has a small streamlined diploid genome (22 Mb haploid) with a high-quality phased genome sequence available to the research community (Craig et al., 2025). Its genetic tractability by homologous recombination makes it a powerful model system in which hypotheses can be tested through reverse genetics (Craig et al., 2025; Dueñas et al., 2025). It can also be cultivated under phototrophic, mixotrophic, and heterotrophic conditions. Under heterotrophic and mixotrophic conditions, the photosynthetic apparatus is degraded, and triacylglycerol accumulates (Matsuka et al.,1966), providing a compelling system for investigating biogenesis and regulation of photosynthesis.

Interestingly, a previous analysis of photosynthesis-related genes in *Auxenochlorella* failed to identify orthologs of either LHCSR or PSBS (Craig, et al 2025). Nevertheless, a PSBS-like sequence (*PSBSL1)* is found in the *Auxenochlorella* genome (Craig et al., 2025). Furthermore, we identified genes encoding both a putative VDE (*VDE1*) and a putative CVDE (*CVDE1*), raising questions about how this alga performs qE and qZ. Here, we combined phylogenetic, molecular, and genetic approaches to characterize NPQ and the *in vivo* functions of genes encoding homologs of PSBS, VDE, and CVDE in the emerging model green alga *Auxenochlorella*.

## Materials and Methods

### AlphaFold

Structural models of *Arabidopsis thaliana,* from hereafter *Arabidopsis,* VDE (AT1G08550), *Chlamydomonas* CVDE (Cre04.g221550.t1.2), and *Synechococcus sp. PCC 7002* CruP (SYNPCC7002_A0043) were generated using the AlphaFold Server (AlphaFold3; Google DeepMind), which implements the AlphaFold neural network architecture for protein structure prediction (Jumper et al., 2021). Transit peptides were predicted using TargetP 2.0 (Almagro Armenteros et al., 2019) and removed to generate a mature protein sequence. Mature protein sequences were submitted to the AlphaFold Server, and the highest-confidence model (ranked by predicted local distance difference test, pLDDT) was used for downstream structural analysis and figure generation.

### Generation of mutants

*Auxenochlorella protothecoides x symbiontica* UTEX 250-A derived from a single colony of *Auxenochlorella protothecoides x symbiontica* UTEX 250 (University of Texas Culture Collection of Algae [UTEX 250]) was used to generate mutants in *VDE1* (UTEX250_A10.26655, UTEX250_B10.26655) and *CVDE1* (UTEX250_A11.34560, UTEX250_B11.34560). The *psbsl1* mutant was generated in the parental strain, *Auxenochlorella* UTEX 250, which was heterozygous at the *PSBSL1* locus (UTEX250_PSBSL1-1 and UTEX250_PSBSL1-2, differing by two amino acids). PSBSL1-1and PSBSL1-2 protein and CDS sequences are listed in supplementary materials. The genome of UTEX 250-A is homozygous for *PSBSL1*; the two alleles UTEX250_A08.18235 and UTEX250_B07.18235 encode the same protein as UTEX-250 PSBSL1. The reference strain used for the *psbsl1* mutant is the appropriate parental strain, UTEX 250.

Gene replacement constructs were designed according to methods described by Craig et al. (2025) (primers listed in supplementary material). All constructs were assembled into a pUC19 backbone using NEBuilder HiFi Assembly (New England Biolab, Ipswich, Massachusetts) and transformed into *Auxenochlorella* UTEX 250-A or UTEX 250 as described by Craig et al., 2025. Mutants disrupted in a single gene (*psbsl1*, *vde1*, and *cvde1*) were generated through two sequential rounds of homologous recombination (one for each chromosome), whereas generation of *vde1 cvde1* required four sequential rounds. In the *psbsl1* mutant, the two *PSBSL1* alleles were replaced with a thiamine biosynthesis (*THIC)* cassette, enabling thiamine prototrophy, and a *nptII* neomycin resistance (Neo^R^) cassette, conferring resistance to 100 μg/mL G418 (Craig et al., 2025). In the *vde1* mutant, the two *VDE1* alleles were replaced with a Neo^R^ cassette and a *THIC* cassette. In the *cvde1* mutant, the two *CVDE1* alleles were replaced with Neo^R^ and *aphVII* hygromycin resistance (Hyg^R^) cassettes (conferring resistance to 300 μg/mL hygromycin B (Craig et al.,2025). The *vde1 cvde1* mutant contained the same marker combinations used in the corresponding single mutants. The double mutant *vde1 cvde1* was generated from the *vde1* (Thi^+^Neo^R^) background using a Hyg^R^ and a second Neo^R^ selection cassette (conferring resistance to 200 μg/mL G418). Thiamine diphosphate is an essential cofactor for 1-deoxy-D-xylulose-5-phosphate synthase, the rate-limiting enzyme of the plastidial methylerythritose 4-phosphate pathway that produces precursors for carotenoid and chlorophyll (phytol) biosynthesis (Lois et al., 2000). Because mutants *vde1* and *vde1 cvde1* were generated by introducing thiamine prototrophy that could affect carotenoid and chlorophyll profiles, a *THIC* control strain (THIC_ref_) was generated by inserting the *THIC* cassette into a neutral site, *THI4* (UTEX250_A10.28900). Schematics of the constructs are shown in Fig. S1. Mutant genotypes were confirmed by colony PCR and Sanger sequencing of genomic DNA (mutant confirmation and sanger sequencing in supplementary material).

### Culture conditions

Cells were grown under white LED lights at 100 µmol photons m^-2^ s^-1^, 25°C, and 200 rpm in an INFORS HT incubator shaker (INFORS HT, Bottmingen, Switzerland) in ACORE media. ACORE (*Auxenochlorella protothecoides*-Optimized Ratio of Elements) medium was based on the CORE (*Chromochloris*-Optimized Ratio of Elements) medium (Jeffers et al. 2025). The ACORE formulation differs from CORE medium in its nitrogen source, with 7.5 mM (NH₄)₂SO₄ replacing 45 mM NaNO₃, and in the addition of 2 µM thiamine-HCl. All other macronutrient, micronutrient, and HEPES concentrations were maintained as described for CORE medium. Growth in low light (LL) was at a light intensity of 100 µmole-photons m^-2^ s^-1^ and in high light (HL) was at 300 µmole-photons m^-2^ s^-1^. Cell density was measured using a Beckman Coulter Multisizer cell counter (Beckman Coulter, Indianapolis, IN, USA).

### NPQ measurement

Cultures were grown to 10 x 10^6^ cells mL^-1^ and 1 mL volumes were pipetted into 12-well plates. Cells were dark-acclimated for 30 minutes before NPQ measurement using an Imaging-PAM M-Series fluorometer (Walz, Germany). The actinic light treatment (300 µmol photons m^-2^ s^-1^) lasted 10 minutes followed by 10 minutes of darkness, with 1190 µmol photons m^-2^ s^-1^ saturation pulses to measure F_m_ and F_m_’. NPQ was calculated as (F_m_ – F_m_’)/F_m_’.

### Identification of PSBS-like, VDE, and CVDE proteins

PSBS-like protein sequences were identified based on sequence similarity to *Auxenochlorella* PSBSL1 by either a jackhmmer search (HMMER v3.41) (Rajkovic et al., 2026) with two iterations against the UniProt database (UniProt Consortium, 2025) (database: Reference Proteomes 2025_01) using default parameters or a BLASTp search against green algal proteomes in the ClusteredNR Database (dated July 27, 2026) using the BLASTp (Altschul et al., 1990) web service at the National Center for Biotechnology Information (NCBI) with default parameters. To differentiate PSBS-like sequences from related proteins sharing the chlorophyll a/b binding protein domain, a sequence similarity network (SSN) generated with full-length sequences was used: proteins more similar to CrPSBS1 (Cre01.g016600), CrPSBS2 (Cre01.g016750), and CrPSBS3 (Cre03.g146147) than to other plant and algal chlorophyll *a*/*b* binding proteins were used for downstream analyses. These sequences were aligned using the MUltiple Sequence Comparison by Log-Expectation (MUSCLE) (Edgar, 2004) alignment tool accessed through the EMBL-EBI Job Dispatcher (Madeira et al., 2024). Jalview v2.11.5.2 (Waterhouse et al., 2009) was used to visualize the resulting alignment, to remove columns with 10% or less occupancy, and to remove land plant sequences with gaps larger than 20 amino acids. The PSBS sequence alignment from Green and Pichersky (1994) was used to divide the aligned sequences into N-terminal and C-terminal regions. The resulting subsequences were used to build a sequence similarity network (SSN) (Oberg et al., 2023) using an alignment score of 15. A SSN built with an alignment score of 20 was used to identify clusters.

VDE protein sequences were identified based on sequence similarity to *Arabidopsis* VDE1 (AT1G08550) and *Chlamydomonas* CVDE1 proteins sequences based on similarity to CVDE (Cre04.g221550) using the methods described for PSBS above using jackhmmer searches and the UniProt database. The resulting sequences were used to build SSNs using the EFI-EST webtool (Oberg et al., 2023) with alignment scores as indicated in the figure legend.

### HL exposure and gene expression analysis by quantitative reverse transcriptase (qRT)-PCR

WT cells were grown to approximately 5 x 10^6^ cells mL^-1^ and transferred to a light intensity of approximately 300 µmol photons m^-2^ s^-1^. Cells were collected at 0,1, 2, and 4 hours for *PSBSL1* and flash frozen in liquid nitrogen. DNA-free total RNA was extracted from frozen algal samples and purified using the Qiagen RNeasy Mini Kit (QIAGEN, Hilden, Germany). RNA concentration and quality were assessed using a NanoDrop spectrophotometer. cDNA was synthesized using the Qiagen Omniscript RT Kit (QIAGEN, Hilden, Germany) following the manufacturer’s instruction using 1 µg of total RNA.

Primer pairs were designed to specifically amplify tubulin (UTEX250_B12.36910.1), *PSBSL1-1*, *PSBSL1-2* and both *PSBSL1* alleles transcripts (primers listed in supplementary material). Primer efficiencies were determined using serial cDNA dilutions (1, 1/3, 1/5, 1/9, and 1/27) and standard curve analysis (primer efficiency listed in supplementary materials). Only primer pairs with amplification efficiencies between 90% and 105% were used. Relative transcript abundance was calculated using the ΔΔCt method (Livak et al.,2001)

### Iso-Seq read alignment to VDE1 and CVDE1 gene models

To assess Iso-Seq reads for the *VDE1* (UTEX250_B10.26655) and *CVDE1* (UTEX250_B11.34560) genes in *Auxenochlorella protothecoides* × *symbiontica* UTEX 250-A (haplotype B-C), genomic sequences for each locus were used as BLASTn (Altschul et al., 1990) queries against the NCBI BLAST web interface (blast.ncbi.nlm.nih.gov), using default search parameters. Each query was searched against a specific PacBio Iso-Seq dataset generated from UTEX 250-A grown autotrophically (SRA experiment SRX27360478, run accession SRR32007620; BioProject PRJNA1211023), by specifying the run accession directly in the BLAST “Sequence Read Archive (SRA)” database field. Reads aligning to each locus were identified from the resulting hit tables, and the number of unique supporting reads was determined by collapsing multiple alignment blocks (representing individual exons) belonging to the same read. Splice-junction positions were inferred from the gaps in query-sequence coordinates between adjacent aligned blocks for each read and compared across all reads mapping to a given locus to assess consistency of the predicted exon-intron structure.

### Dithiothreitol (DTT) treatment

Wild-type (WT) cells were grown to a density of approximately 10 x 10^6^ cells mL^-1^. Freshly prepared 1 M DTT was added to cultures to final concentrations of 0, 5, 10, 20, or 30 mM during a 30-minute dark acclimation period. NPQ was partially inhibited by 5 mM DTT, and additional DTT concentrations did not further suppress NPQ.

### TCSPC

Time-Correlated Single-Photon Counting (TCSPC) measurements were performed to measure fluorescence lifetimes of chlorophylls following an excitation pulse under different actinic light conditions. Samples were grown at 100 µmol photons m^-2^ s^-1^, 25°C, and 250 rpm until OD_700_ reached 1.15. Optical density was checked via the Molecular Devices SpectraMax ABS. A Coherent Verdi G10 diode laser at 532 nm was used to pump a Coherent Mira 900f Ti:sapp oscillator operating at 76 MHz, generating pulses with a fundamental frequency of 808 nm. This beam was frequency-doubled to 404 nm using a beta-barium borate crystal to selectively excite the Q_x_ band of Chl *a*. Using a beamsplitter, part of the beam was directed to a Becker-Hickl sync photodiode (PHD-400) to provide synchronization references to the Becker-Hickl TCSPC card (PSC-150). The remaining beam was directed at the surface of a cuvette containing an *Auxenochlorella* sample in detergent at an angle of approximately 70^°^.

For WT and *cvde1* samples, data were collected using a laser power of 1.0 mW. For the THIC_ref_, *vde1* and *vde1 cvde1* samples, data were collected using a laser power of 0.6 mW.

Before each TCSPC measurement, samples were dark acclimated for 80 minutes to ensure NPQ processes were deactivated. Data were collected in snapshots taken every 15 s, which used a laser excitation and detector integration time of 1 s. The first three snapshots were taken in the dark to measure baseline fluorescence lifetimes for each sample, after which the actinic light sequence began. We used a sequence of 20 minutes HL, 20 minutes dark, 20 minutes HL, 20 minutes dark. HL was at a light intensity of 300 μmol photons m^-2^ s^-1^ as in other experiments and dark was complete darkness.

Chl *a* Q_y_ band fluorescence emission was detected at 680 nm using a HORIBA Jobin-Yvon monochromator (H-20) and a Hamamatsu R3809U microchannel plate-photomultiplier tube (MCP-PMT) detector. The instrument response function (IRF) for our system had a full-width half-maximum of 37 ps, sufficient to resolve sub-ns fluorescence lifetime changes. LabVIEW codes were used to control shutters for the excitation beam, turn the actinic light on and off, and start detection based on input sequences.

Each collected fluorescence decay profile was convoluted with the IRF:

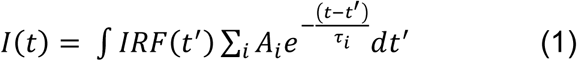

This curve was fit with a biexponential decay function, from which an amplitude-weighted average fluorescence lifetime was calculated:

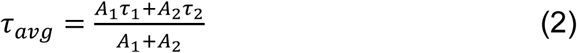

The average fluorescence lifetimes were then used to calculate a dimensionless NPQ value, expressed as a change in fluorescence lifetime relative to the averaged lifetime of the first three dark snapshots:

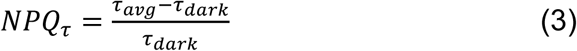

### HPLC

WT cultures were grown to approximately 5 x 10^6^ cells mL^-1^. Cultures were dark acclimated overnight (∼12 hours) before exposure to approximately 300 µmol photons m^-2^ s^-1^ light. Samples were collected 0, 15, 30, and 60 minutes after onset of illumination, flash frozen in liquid nitrogen, and stored until extraction. Samples were extracted according to Jeffers et al., 2025 and analyzed by HPLC. Pigments were separated on a C30 reversed-phase YMC Carotenoid column (250 × 4.6 mm I.D., 5 µm particle size; column volume 4.2 mL) (YMC Co., Ltd., Kyoto, Japan) following the method described by Gupta et al., 2015.

Following fluorescence measurements, *Nicotiana benthamiana* (hereafter *Nicotiana*), leaf punches were dark acclimated for an additional 30 minutes and exposed to 1000 µmol photons m^-2^ s^-1^ light for 30 minutes. Samples were immediately flash frozen in liquid nitrogen and stored until pigment extraction. Pigments were extracted from frozen tissue on ice and in the dark using ethanol and analyzed by HPLC using the method previously described.

### Statistical Tests and Data Visualization

Claude (Anthropic) was used to assist with the development and design of code for data visualization of fluorescence, HPLC, and qPCR data, as well as code for performing two-way ANOVA and Tukey’s HSD post hoc tests. All AI-assisted code was reviewed, independently checked, and validated by the authors for accuracy and appropriate implementation prior to use in the analysis and presentation of the results. Tukey HSD was run separately at each timepoint using replicate-level values. For each reference– mutant pair and pigment, a two-factor model was fit with Strain, Time, and Strain × Time. Reported sums of squares are partial/Type III-like tests from nested-model comparisons.

### Transient Expression

Transit peptides and mature proteins were predicted using TargetP (https://services.healthtech.dtu.dk/services/TargetP-2.0/). The predicted mature *Auxenochlorella* VDE1 protein sequence was fused to the *Arabidopsis* VDE1 transit peptide (*At*VDEtp). The resulting *At*VDEtp-*Apxs*VDE1 construct, and the native *Auxenochlorella* CVDE1 sequence were codon optimized for expression in *Nicotiana* using Genscript codon optimization tool (https://www.genscript.com/tools/gensmart-codon-optimization).

The codon-optimized constructs, together with native *Nicotiana* (*Nb*VDE) and a GUS control, were independently cloned into a pUBQ-mScarlet vector using HiFi assembly. Plasmids were transformed into XL1-Blue competent cells and verified by whole-plasmid sequencing. Verified plasmids were subsequently transformed into *Agrobacterium tumefaciens* strain GV3101.

*Agrobacterium* strains were streaked onto LB agar containing 50 µg/mL kanamycin, 30 µg/mL gentamicin, and 10 µg/mL rifampin and incubated at 28°C for 48 hours. Individual colonies were used to inoculate duplicate 5 mL LB cultures containing the same antibiotics. Cells were harvested by centrifugation and resuspended in infiltration medium (97 mL H₂O, 1 mL 1 M MgCl₂, 2 mL 500 mM MES pH 5.6, and 150 µL 100 mM acetosyringone). Cultures were incubated in the dark at room temperature for 3 hours before being adjusted to an OD_600_ of 0.75.

WT and *npq1 Nicotiana* plants (Lee et al., 2024) were grown for approximately 2.5 weeks in a temperature-controlled growth chamber with 14 hour days at 150 μmol photons m^-2^ s^-1^ light, temperature was 27°C during the day and 23°C at night. Emerging shoots and visibly damaged leaves were removed before infiltration. Approximately 1 mL of each *Agrobacterium* suspension was infiltrated into separate regions of healthy young leaves.

Leaf punches were collected 72 hours post-infiltration and dark acclimated for 30 minutes before NPQ measurements. Relative mScarlet fluorescence was quantified using a Tecan Infinite M1000 Pro plate reader (Tecan Group Ltd., Männedorf, Switzerland) immediately following NPQ measurements. Fluorescence measurements were performed using excitation at 570 nm and emission at 600 nm with excitation and emission bandwidths of 8 nm and a gain setting of 75.

## Results

### Wild-type *Auxenochlorella* lacks rapidly reversible qE and a functional PSBS

The absence of *LHCSR* and *PSBS* in the *Auxenochlorella* genome raised the question whether *PSBSL1* has an essential function for NPQ in this alga. To test this, we generated *psbsl1* mutants by knocking out each allele sequentially, generating single-allele knockout (*psbsl1-1/PSBSL1-2*) and double-allele knockout (*psbsl1*) mutants (Fig. S1). NPQ was measured in comparison to the wild type (WT) to determine whether the NPQ capacity was affected by the light environment to which the cells have acclimated, NPQ was measured in cells grown in low light (LL; 100 µmol photons m^-2^ s^-1^) and high light (HL; 300 µmol photons m^-2^ s^-1^).

Both the *psbsl1-1/PSBSL1-2* and *psbsl1-1/psbsl1-2* showed no significant difference in NPQ compared to WT, which exhibited a relatively low induction of NPQ (∼1), which relaxed slowly in the dark (Fig. 2a). HL-grown cells had marginally increased NPQ with slower relaxation in the dark compared to their LL-grown counterparts. These results showed that *Auxenochlorella* lacks rapidly reversible qE and exhibits a low level of slowly reversible NPQ that is unaffected by the deletion of *PSBSL1*, strongly suggesting that this gene does not encode a PSBS that functions in qE.

**Figure 2.**
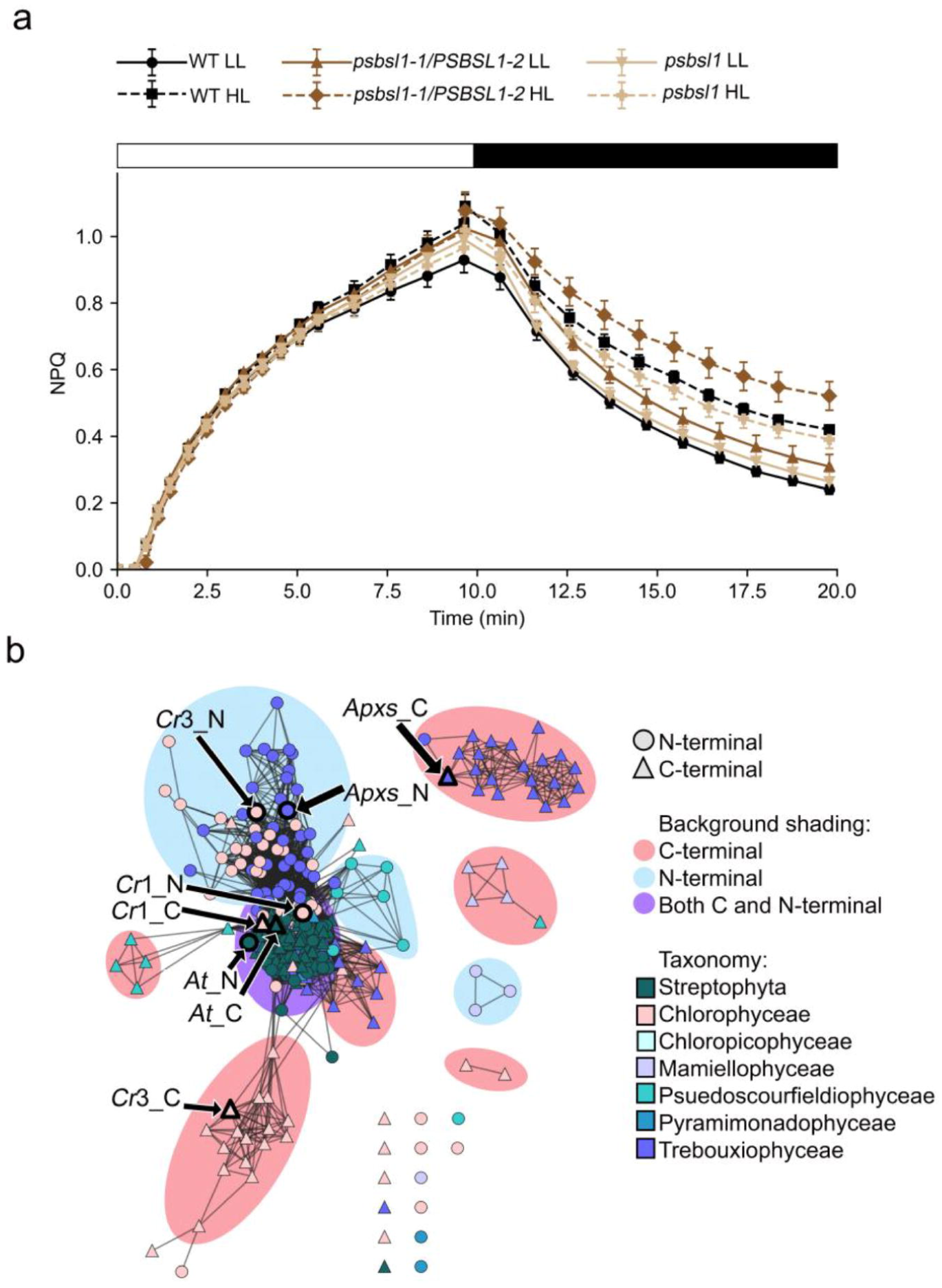
Role of PSBSL1 protein in *Auxenochlorella* NPQ. **a)** NPQ of WT (*Auxenochlorella* UTEX-250) *psbsl1-1/PSBSL1-2,* and *psbsl1-1/psbsl1-2* grown in LL and HL. White bar represents illumination with 300 µmol photons m^-2^ s^-1^ actinic light, black bar represents dark (with low measuring light) in the NPQ script. Values represent averages and standard error shown as error bars (n=3). **b)** SSN of PSBS and PSBS-like proteins. Nodes are colored by taxonomy according to the color key, and node shape indicates whether the node represents an N-terminal pair of helices (circle) or C-terminal pair of helices (triangle). Background shading indicates whether the cluster of nodes are predominately N-terminal (blue), C-terminal (pink), or a mix of N- and C-terminal pair of helices (purple). The location of sequences corresponding to *Arabidopsis* PSBS (*At*), *Auxenochlorella* PSBSL1 (*Apxs*), *Chlamydomonas* PSBS1/2 (*Cr*1) and PSBS3 (*Cr*3) are indicated with an arrow. Gene models used; *At*PSBS (AT1G44575), *Cr*PSBS1 (Cre01.g016600,Cre01.g016750),*Cr*PSBS3 (Cre03.g146147) and *Apxs*PSBSL1 (B07.18235, A08.18235). Proteins not associated with a defined cluster are shown outside the shaded cluster regions.

In *Chlamydomonas,* the gene expression of *LHCSR* and *PSBS* is strongly induced by HL (Mettler et al., 2014). In contrast, we found that the *Auxenochlorella PSBSL1* gene was repressed when LL-grown cells were transferred to HL (Fig. S2a), a typical pattern observed for *LHC* genes, such as *LHCBM* genes (Teramoto et al., 2002; Durnford et al., 2003; Elrad and Grossman, 2004).

To understand the relationships among the proteins similar to PSBS within the LHC protein family, we surveyed plant and green algal LHC proteins with four transmembrane helices, starting with PSBS from *Arabidopsis* and *Chlamydomonas.* Sequence similarity network (SSN) analysis (see methods) of the helices in *Arabidopsis* PSBS and *Chlamydomonas* PSBS showed that the first and second (N-terminal) helices are homologous to the third and fourth (C-terminal) helices, which was not the case for many other proteins similar to PSBS (Fig. 2b). Furthermore, other known qE-mediating PSBSs exhibited the same pattern of N-terminal helix pairs being homologous to C-terminal helix pairs (Fig. S2b and c), whereas the N-terminal and C-terminal helix pairs in PSBSL1 and a *Chlamydomonas* protein with limited similarity to PSBS (PSBS3) were divergent (Fig. 2b). Based on these results, we consider the *Auxenochlorella* gene to encode a PSBS-like protein rather than a typical PSBS. Altogether, we conclude that *Auxenochlorella* encodes neither a PSBS that functions in qE nor an LHCSR in its genome, which explains the lack of qE.

### *Auxenochlorella* has candidate genes for both plant-type VDE and CVDE

Given the small total NPQ and its slow induction and relaxation kinetics (Fig. 2a), we hypothesized that NPQ in *Auxenochlorella* mainly consists of qZ. Therefore, we searched for homologs of *Arabidopsis VDE1* and *Chlamydomonas CVDE1* in the *Auxenochlorella* genome and found candidates for both genes.

The identified *Auxenochlorella* VDE1 clustered with VDE proteins from other green algae and land plants along with those from Stramenopila, Alveolata, and Rhizaria (SAR), supporting its annotation as a plant-like VDE (Fig. 3a). Alignment of the *Auxenochlorella* VDE1 sequence with the well-characterized *Spinacia oleracea* and *Arabidopsis* VDE sequences (Fig. S3a) confirmed the presence of a distinct cysteine-rich N-terminal domain, a lipocalin domain, and a C-terminal domain found previously in *S. oleracea* VDE, including residues that are essential for enzymatic activity (Hieber et al., 2000, 2002; Saga et al., 2010; Hallin et al., 2015). Multiple protonatable residues were also conserved, although several acidic residues displayed D/E substitutions. These substitutions are unlikely to substantially alter protonation behavior, due to the similar chemical properties and charges of aspartate and glutamate (Goldstein and Pollock, 2016). Sequence comparisons suggested that *Auxenochlorella* VDE1 has properties similar to functional plant-like VDEs and may represent a functional VDE.

**Figure 3.**
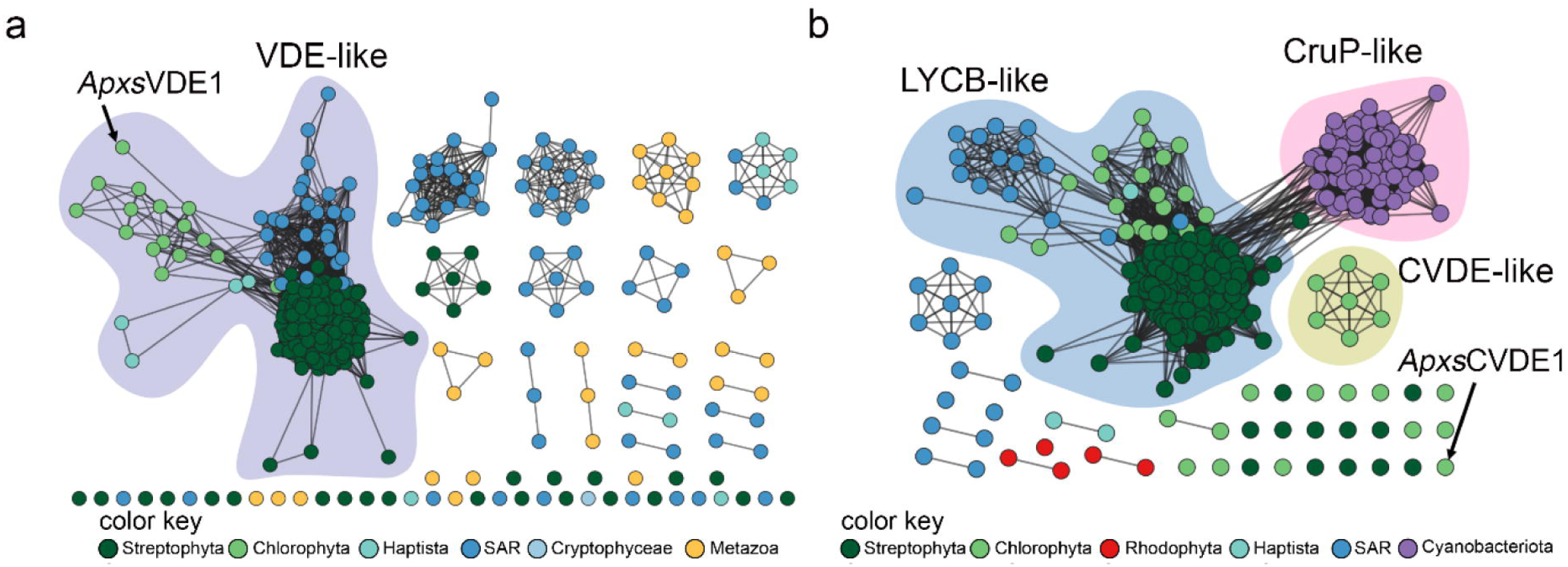
Characterization of *Auxenochlorella* VDE1 and CVDE1. SSN of proteins similar to **a)** VDE and **b)** CVDE. Nodes are colored by taxonomy according to the color key. Background shading and accompanied label indicate protein clusters for VDE-like proteins (purple), LYCB-like (blue), CruP-like (pink) and CVDE (gold). The location of sequences corresponding to *Auxenochlorella* are indicated with an arrow. Proteins not associated with a defined cluster are shown outside the shaded cluster regions.

Due to their paralogous relationship, algal CruP and CVDE proteins formed closely related clusters, with the putative *Auxenochlorella* CVDE1 not grouping with the CruP-like or CVDE clusters (Fig. 3b). A protein alignment of *Auxenochlorella* CVDE1 with the protein most similar to *Synechococcus sp. PCC 7002* CruP in *Chlamydomonas,* hereafter called *Chlamydomonas* CruP (Cre06.g264250), and *Chlamydomonas* CVDE1 were generated to further confirm the identity of the putative *Auxenochlorella* CVDE1 (Fig. S3b). The N-terminal regions of *Chlamydomonas* CruP, CVDE1, and *Auxenochlorella* CVDE1 had predicted FAD-binding oxidoreductase domains. *Chlamydomonas* CruP and CVDE1 contain distinct FAD oxidoreductase-binding domains, DAO and UbiH, respectively, and *Auxenochlorella* CVDE1 showed conservation of the UbiH domain seen in *Chlamydomonas* CVDE1 (Supplementary Fig. S3b). Specific residues of importance for CVDE enzymatic activity have not yet been identified, preventing further residue-level annotation. The shared UbiH domain between *Chlamydomonas* CVDE1 and *Auxenochlorella* CVDE1 suggests that *Auxenochlorella* CVDE1 may also be a CVDE.

Iso-Seq reads from *Auxenochlorella* grown under autotrophic conditions (Craig et al., 2025 dataset; SRA: SRR32007620) were used as a rough approximation of the relative transcript abundance of *VDE1* and *CVDE1* under basal autotrophic conditions, given the differing read depths and non-quantitative nature of Iso-Seq library preparation. A total of 425 Iso-Seq reads aligned to the *VDE1* locus, of which 96.5% (410/425) of reads were shorter, beginning partway through the gene body and lacking the predicted transit peptide, N-terminal domain and part of the lipocalin domain. Notably, only 3.5% (15/425) of reads extended far enough in the 5’ direction to capture the full-length transcript. In contrast, only 12 Iso-Seq reads aligned to the *CVDE1* locus, of which 75% (9/12) supported a shorter transcript containing three exons. The remaining 25% (3/12) of *CVDE1* reads showed exon alignment patterns that did not match the boundaries seen in the other 9 reads. Each divergent pattern was observed in only a single read, and we cannot distinguish whether this reflects genuine alternative splicing or an alignment artifact (e.g., from a sequencing error or ambiguous sequence near the splice site).

We further explored the contribution of VDE1 to NPQ in *Auxenochlorella* by treating cells with DTT. The activity of VDE, but not CVDE, is inhibited by DTT (Li et al., 2016), and this allowed us to investigate the contribution of VDE1 to NPQ induction. Cells were treated with varying concentrations of DTT prior to NPQ measurements. DTT treatment decreased maximal NPQ by approximately 50% compared to the control (Fig. 4a). To examine whether the decrease in NPQ was accompanied by inhibition of VDE1 activity, we examined the accumulation of VAZ cycle xanthophylls (Vio, Anthera, and Zea). Previous studies have shown that the rise and stabilization of Zea and qZ is most observable within the timeframe of 0 to 30 minutes (Nilkens et al., 2010; Kress & Jahns, 2017). Thus, we monitored VAZ cycle xanthophylls during 30 minutes of HL exposure of cells with or without DTT treatment. Additionally, we examined the accumulation of the carotenoid lutein, which has been shown to contribute substantially to NPQ in *Chlamydomonas* independently of xanthophyll-cycle pigments (Niyogi et al., 1997), to assess whether loss of VDE/CVDE function is accompanied by compensatory changes in this parallel photoprotective pigment pool. In non-treated control samples, the VAZ cycle pigments remain relatively stable over time and do not exhibit an inverse pattern between Zea and Vio (Fig. 4b-d). However, the Zea and Anthera from DTT-treated samples remained lower than non-treated control samples over the time course, and lower total VAZ pool and de-epoxidation states (DES) were observed compared to the non-treated control, all of which suggested that VDE activity was indeed inhibited by DTT (Fig. 4b-f). Although we cannot rule out DTT inhibition of other enzymes, the ∼50% reduction in NPQ seen in 5 mM DTT-treated cells was consistent with the hypothesis that qZ is a major component of NPQ in *Auxenochlorella*.

**Figure 4.**
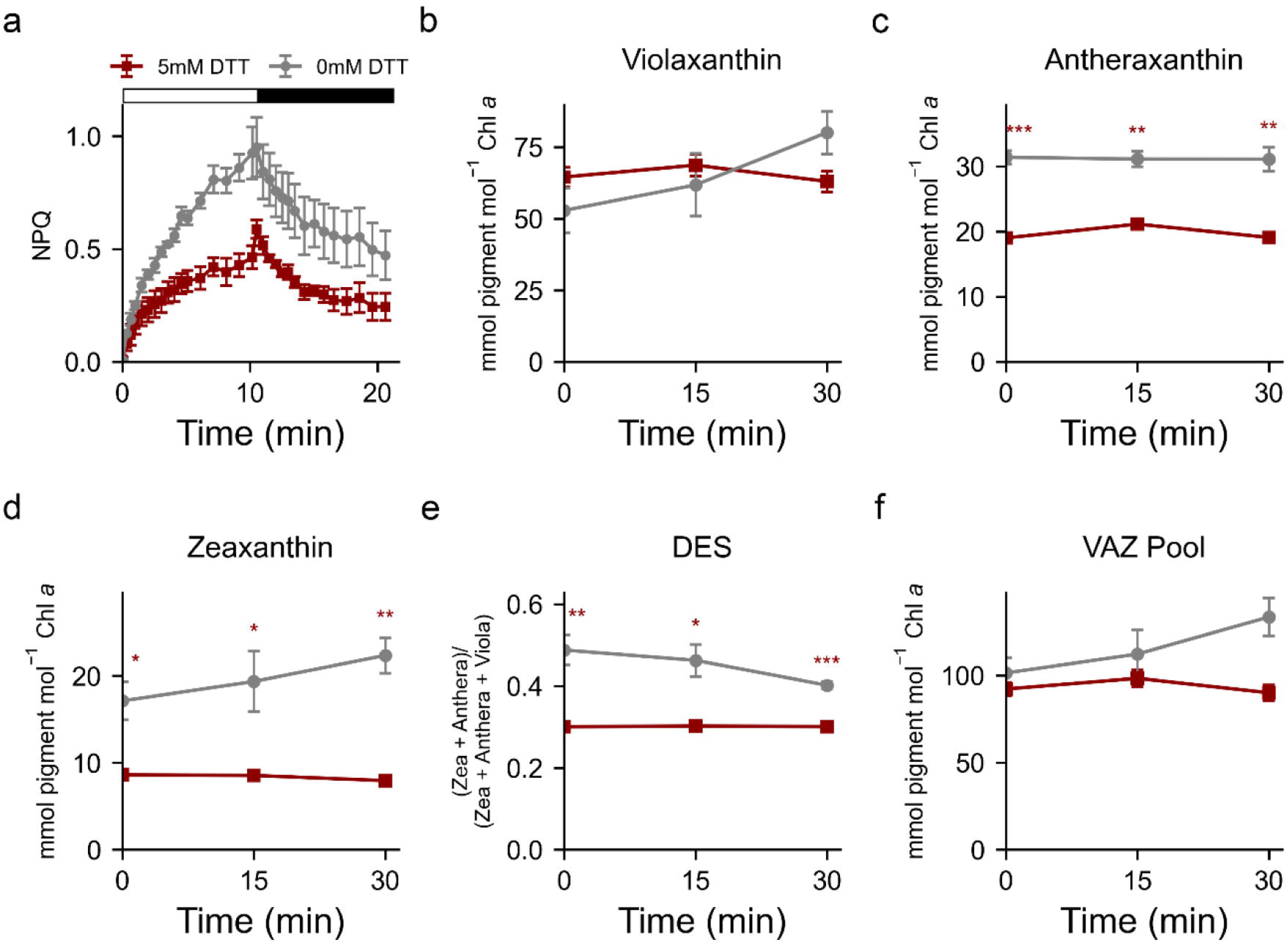
Effects of DTT on *Auxenochlorella* NPQ and xanthophyll cycle pigments. **a)** NPQ of WT *Auxenochlorella* treated with 0 or 5 mM DTT. White bar represents illumination with 300 µmol photons m^-2^ s^-1^ actinic light, black bar represents dark (with low measuring light). **(b-f)** Pigment profiles of DTT treated samples compared to reference strain. **b)** Violaxanthin, **c)** Antheraxanthin, **d)** Zeaxanthin, **e)** DES, **f)** VAZ pool. All pigment concentration was normalized to Chl *a* content in the same sample. Values represent averages and standard error shown as error bars (n=3). Significant differences between reference strain and DTT treatment are indicated by asterisks (Tukey test). Significance levels for Tukey test and Two-way ANOVA: no asterisk = no significance, (*) = P < 0.05, (**) = P < 0.01, (***) = P <0.001.Two-way ANOVA strain effects: antheraxanthin, *F*(1,20) = 38.63, *p* = 0.0167 (*); zeaxanthin, *F*(1,20) = 8.92, *p* = 0.0073 (**); DES, *F*(1,20) = 21.45, *p* = 0.0016 (***).

### *Auxenochlorella* VDE1 and CVDE1 do not contribute to canonical qZ

To investigate further the *in vivo* functions of *Auxenochlorella* VDE1 and *Auxenochlorella* CVDE1, we generated *vde1*, *cvde1*, and *vde1 cvde1* mutants. The *vde1* mutant exhibited approximately 25% decrease in maximal NPQ relative to the reference strain (Fig. 5a). In contrast, the *cvde*1 mutant had a smaller (∼7%) decrease in maximal NPQ compared to WT (Fig. 5b). It is possible that when VDE1 was knocked out, CVDE1 was compensating, and vice-versa. However, the disruption of both genes in *vde1 cvde1* resulted in an increase in maximal NPQ (Fig. 5a) rather than a decrease as expected if compensation was occurring in the single gene mutants, possibly reflecting increased photoinhibition (qI). Overall, these PAM NPQ measurements indicate that VDE contributes more to NPQ than CVDE, which has a minor role. However, the deletion of both *CVDE* and *VDE* renders the alga more photosensitive, suggesting that the two enzymes both contribute to pigment-mediated photoprotection.

**Figure 5.**
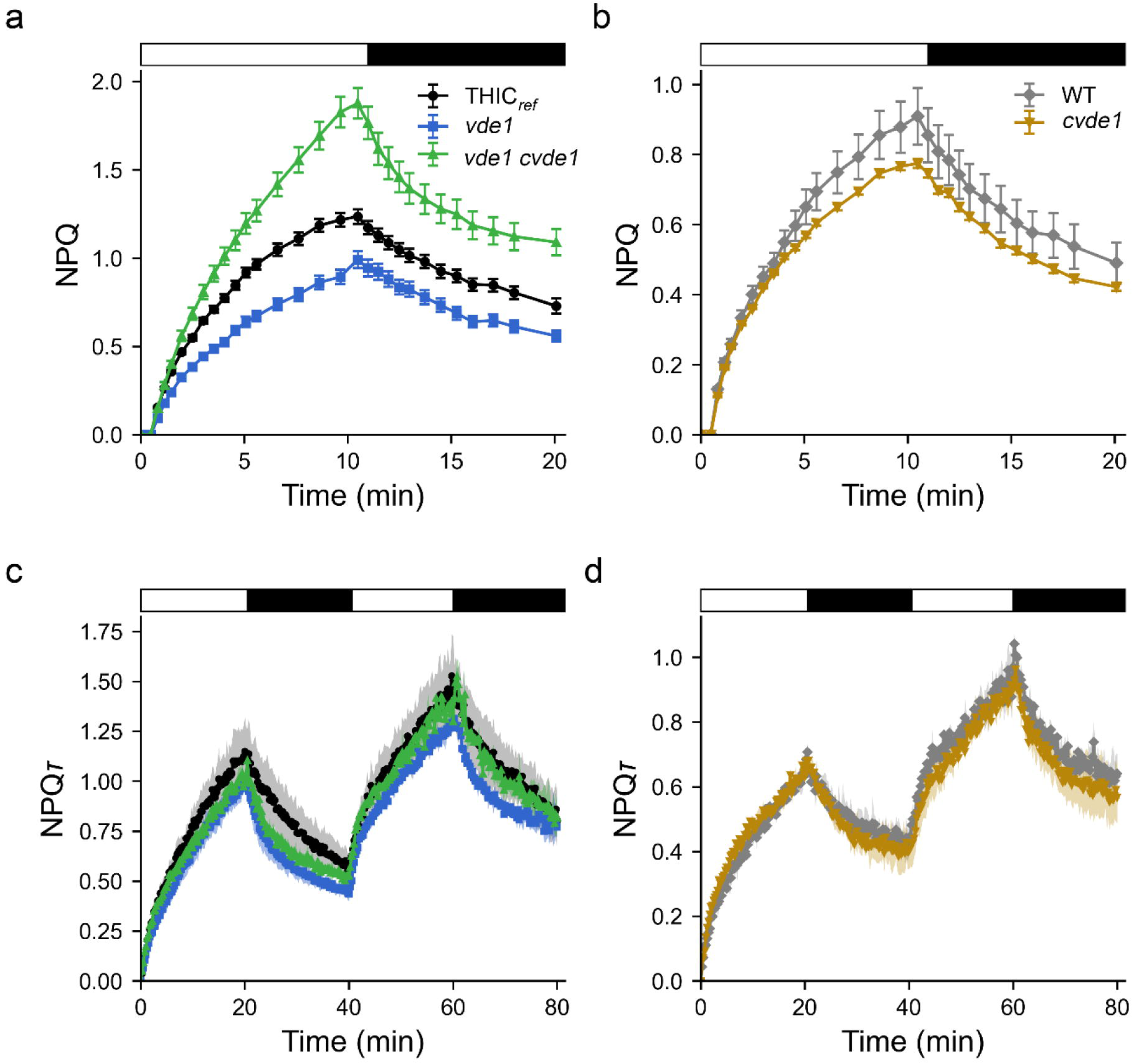
NPQ measured by PAM and NPQτ measured by TCSPC for c*vde1, vde1*, and *vde1* c*vde1* mutants. NPQ measured with PAM for **a)** *vde1, vde1 cvde1,* and reference strain (THIC_ref_) **b)** *cvde1* and WT. Values represent averages (n=4) and standard error shown as error bars (n=4). NPQ τ obtained from TCSPC measurements for **c)** *vde1, vde1 cvde1,* and reference strain (THIC_ref_), for **d)** *cvde1* and WT. Values represent averages and standard error shown as error bars (n=3). White bar represents illumination with 300 µmol photons m^-2^ s^-1^ actinic light, black bar represents dark phase in the NPQ script (with low measuring light).

To further characterize chlorophyll fluorescence dynamics, time-correlated single-photon counting (TCSPC) was performed on the *vde1*, *cvde1*, and *vde1 cvde1* mutants. While the PAM fluorometer measures changes in total fluorescence yield, TCSPC measures changes in chlorophyll fluorescence decay kinetics. This provides a more specific reporter of NPQ dynamics independent of other sources of total fluorescence losses that may affect measurements by PAM, including pigment bleaching (Zaks et al., 2013). The *vde1* mutant exhibited a small decrease in overall NPQτ amplitude relative to the reference strain (Fig. 5c), consistent with the PAM measurement of NPQ (Fig. 5a). The *vde1 cvde1* mutant displayed slightly decreased NPQτ compared to the reference in the first light and dark period, but NPQτ was more similar in amplitude to the reference in the second light and dark period (Fig. 5c). This was in contrast to the PAM measurement, which showed higher NPQ throughout the light and dark phases (Fig. 5a). Contrastingly, we observed that *cvde1* did not exhibit significant changes in NPQτ compared to WT in either the first or second light period, indicating that the excited state lifetimes of pigments involved in quenching were unchanged (Fig. 5d). The differences observed between the PAM and TCSPC of *vde1 cvde1* suggest that the apparent increase in NPQ in *vde1 cvde1* measured by PAM was not due to quenching of the chlorophyll excited-state lifetime. One possible explanation is that pigment bleaching led to inflation of the PAM-based NPQ signal. However, measurement by HPLC did not reveal changes in pigment content consistent with this explanation, as discusssed below, suggesting that the discrepancy reflects other changes that are independent of chlorophyll excited-state quenching.

To understand the effects of *VDE1* and *CVDE1* disruption on carotenoid accumulation and dynamics, we examined pigment profiles of the mutants (Fig. 6 and 7). Cells were grown in LL, transferred to HL and samples were collected at 0, 15, and 30 minutes after transfer. In both reference strains WT and THIC_ref_ (see methods), Zea levels remained relatively unchanged throughout the HL treatment and moreover, it did not exhibit inverse correlation with Vio, a pattern typically associated with qZ and operation of the VAZ cycle (Fig. 6b, c and 7b, c). The *cvde1* mutant exhibited no changes in pigment dynamics and composition compared to the reference strain (Fig. 6a-f). Neither the *vde1* nor the *vde1 cvde1* mutant exhibited changes in VAZ pigment dynamics and VAZ pool size compared to the reference strains but, VAZ pigment composition was altered (Fig.7a-d). The *vde1* and *vde1 cvde1* mutants had increased Anthera and lutein relative to the reference, and the *vde1 cvde1* Vio pool before the HL transfer seemed to be smaller compared to the reference strain (Fig. 7a, b,e, f). The higher Vio and lower Anthera in *vde1 cvde1* was reflected in a higher DES compared to the reference strain (Fig. 7e and f). Overall, we did not observe a complete loss of Zea or Anthera pools in any of the three mutant strains.

**Figure 6.**
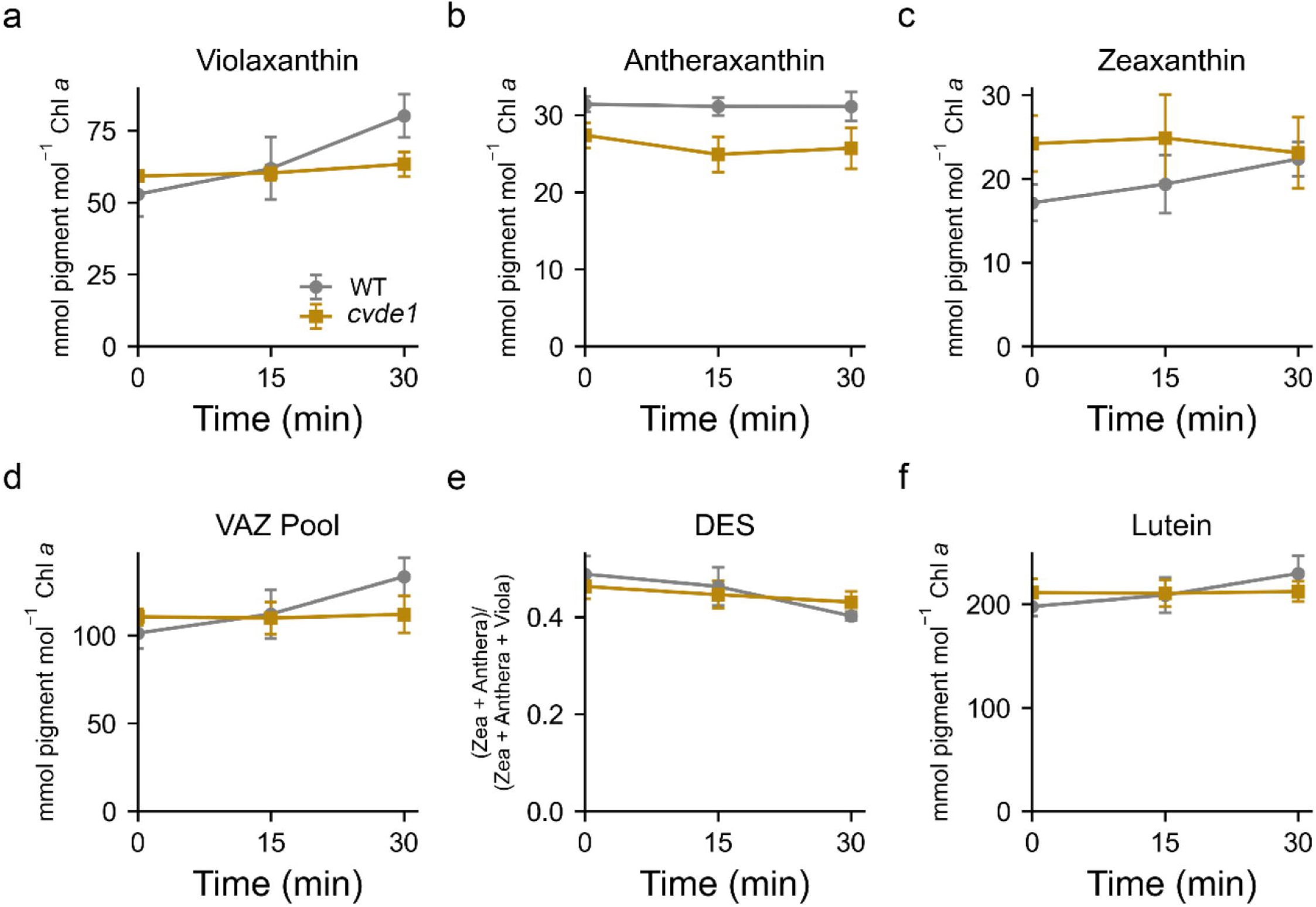
Pigment profile of the c*vde1* mutant compared to WT. **a)** Violaxanthin, **b)** Antheraxanthin, **c)** Zeaxanthin, **d)** Total VAZ pool, **e)** DES, **f)** Lutein. Pigment concentration was normalized to Chl *a* content in the same sample. Values are average of biological replicates and error bars represent standard error (n=4). Significant differences between reference strain and mutants are indicated by asterisks (Tukey test). Significance levels: ns or no asterisk = no significance.

**Figure 7.**
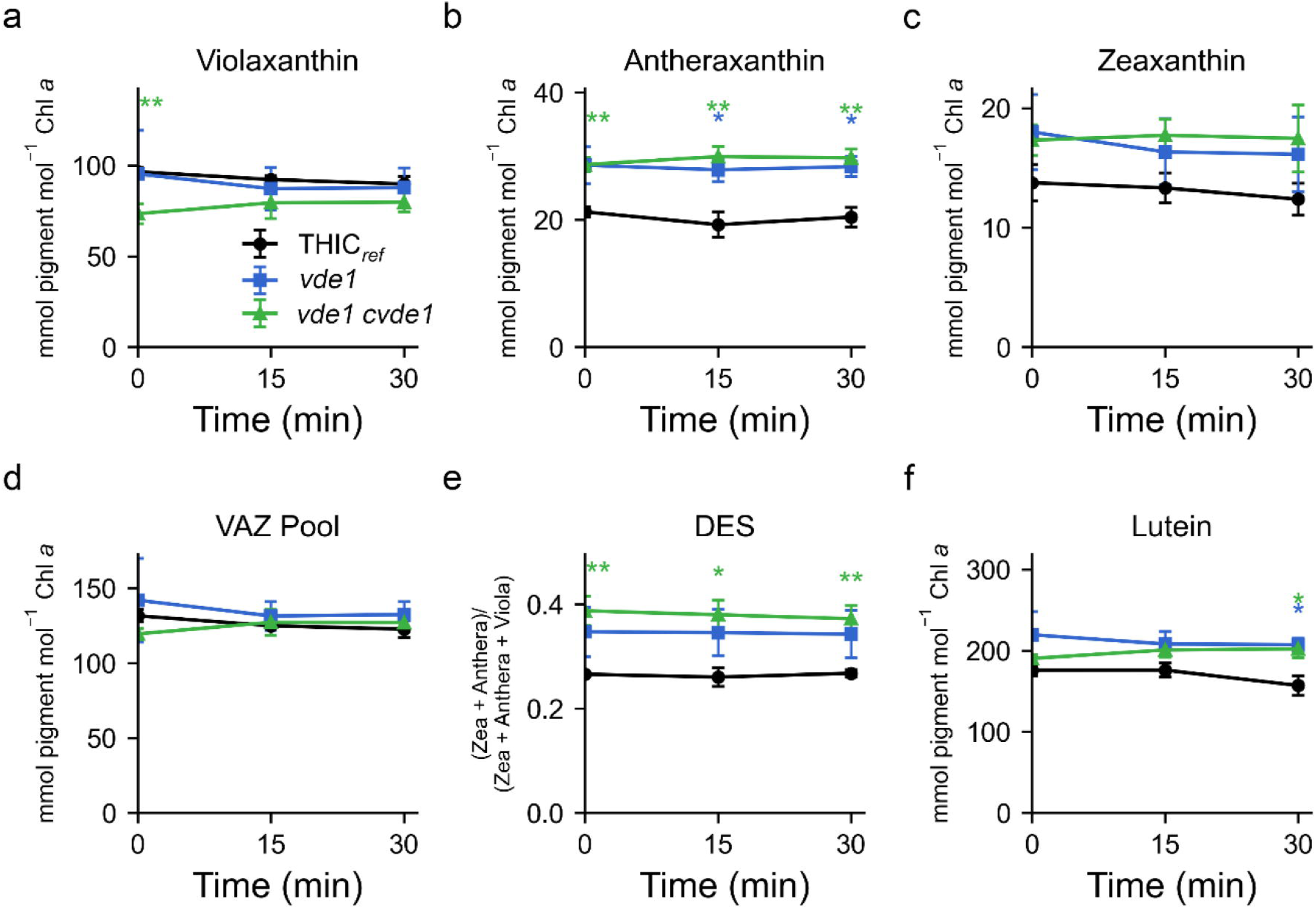
Pigment profile of *vde1* and *vde1* c*vde1* mutants compared to THIC_ref_. a) Violaxanthin, **b)** Antheraxanthin, **c)** Zeaxanthin, **d)** Total VAZ pool, **e)** DES, **f)** Lutein. Pigment concentration was normalized to Chl *a* content in the same sample. Values are average of biological replicates and error bars represent standard error (n=4). Significant differences between reference strain and mutants are indicated by asterisks (Tukey test). Significance levels for Tukey and Two-way ANOVA: ns or no asterisk = no significance, (*) = P < 0.05, (**) = P < 0.0. Two-way ANOVA strain effects: *vde1* — antheraxanthin, *F*(1,24) = 8.92, *p* = 0.0064 (**); lutein, *F*(1,24) = 4.62, *p* = 0.0418 (*). *vde1 cvde1* — antheraxanthin, *F*(1,24) = 10.97, *p* = 0.0029 (**)

### *Auxenochlorella* VDE1 and CVDE1 are active enzymes when heterologously expressed

The lack of canonical qZ-associated VAZ pigment dynamics and the relatively small changes in NPQ observed in the mutants prompted us to investigate if *Auxenochlorella* VDE1 and *Auxenochlorella* CVDE1 had functional de-epoxidase activity. We directly tested this by transiently expressing *Auxenochlorella VDE1* and C*VDE1* in the *Nicotiana npq1* mutant (Lee et al., 2024). Previous work demonstrated that functional green algal CVDE and plant VDE can complement the VDE-deficient *Arabidopsis npq1* mutant and restore Zea production (Li et al., 2016; Sun et al., 2019; Lou et al., 2022). WT and *npq1 Nicotiana* leaves were infiltrated with *Agrobacterium tumefaciens* strains carrying the native *Nicotiana* VDE (*Nb*VDE), *Auxenochlorella* CVDE1 (*Apxs*CVDE1), *Auxenochlorella* VDE1 fused to the *Arabidopsis* VDE1 transit peptide (*At*VDEtp-*Apxs*VDE1), and an mScarlet-pUBQ-GUS control construct (WT-mScarlet and *npq1*-mScarlet) (Fig. 8a). The mScarlet control had minimal effects on NPQ kinetics in both WT and *npq1 Nicotiana* plants (Fig. S4a and b). Following exposure to 30 minutes of HL, mScarlet-infiltrated *npq1* plants contained no detectable levels of Zea and Anthera, as did the *npq1* mutant (Fig. S4c). In contrast, WT plants, mScarlet-infiltrated WT plants, *Apxs*CVDE1 complemented *npq1* plants, and *At*VDEtp-*Apxs*VDE1-complemented *npq1* plants all accumulated Zea and Anthera following HL treatment (Fig. 8b and Fig. S3d). Furthermore, *npq1* plants with *Apxs*CVDE, *Nb*VDE and *At*VDEtp-*Apxs*VDE1 all showed restored NPQ kinetics that were higher in magnitude and faster in relaxation of NPQ compared to WT-mScarlet plants (Fig. 8c), which may be due to the overexpression and overaccumulation of the protein. These results show that *Auxenochlorella* VDE1 and CVDE1 encode function de-epoxidases that can complement *Nicotiana npq1*.

**Figure 8.**
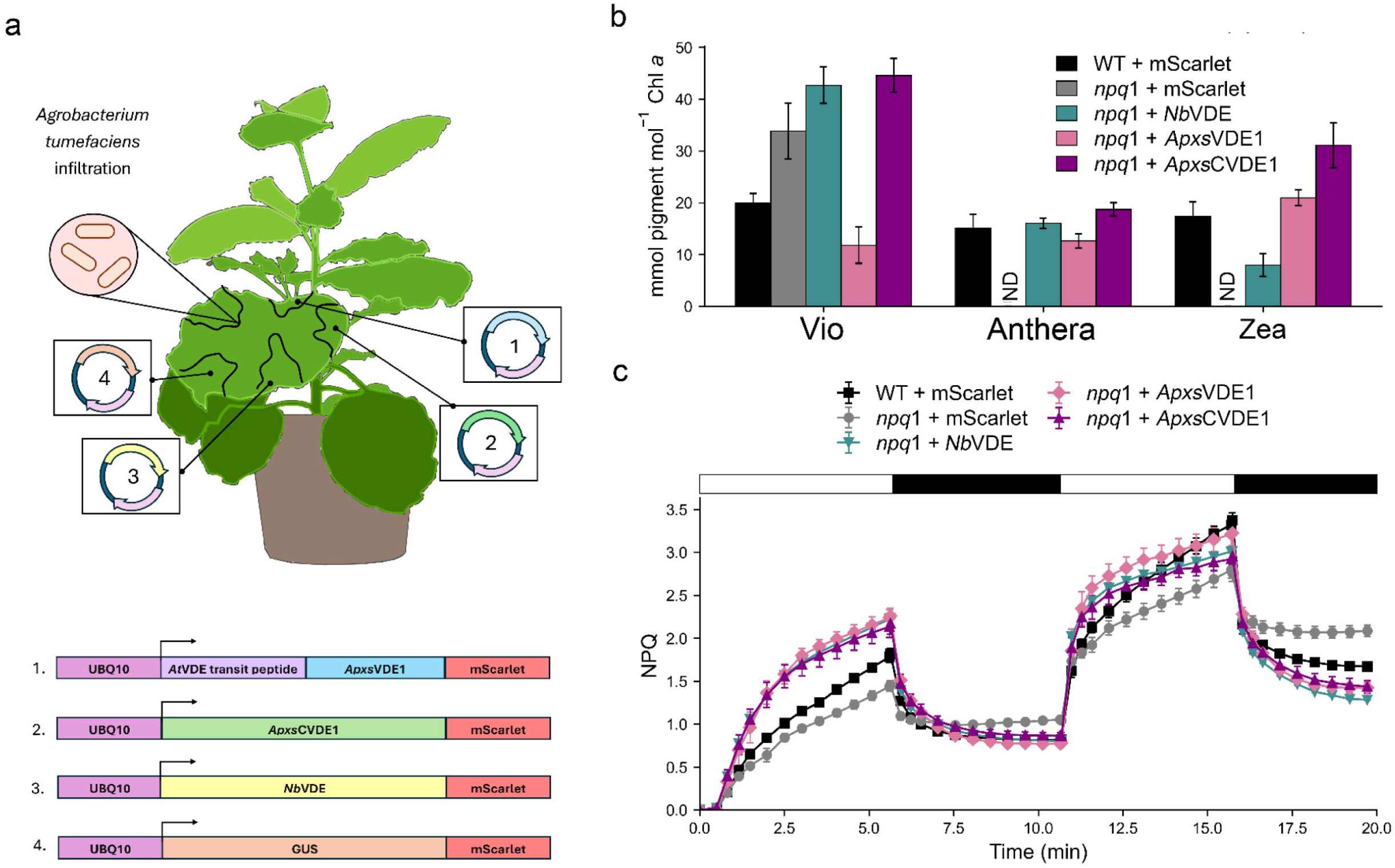
NPQ and pigment profile of *Nicotiana* leaves transiently expressing *Auxenochlorella* VDE1 and CVDE1. **a)** Illustration of *Nicotiana* transient expression and constructs (1,2,3,4). The four constructs were infiltrated into separate regions of the same leaf. NPQ and pigment extraction were performed on the same leaf punches. **b)** Xanthophyll cycle pigments after 30 minutes of illumination with 1000 µmol photons m^-2^ s^-1^ actinic light and, **c)** NPQ measured from WT leaves expressing empty vector (WT + mScarlet), *npq1* leaves expressing empty vector (*npq1* + mScarlet), *npq1-Nb*VDE (*npq1* + *Nb*VDE), *npq1-At*VDEtp-*Apxs*VDE1 (*npq1*+ *Apxs*VDE1) *npq1*-*Apxs*CVDE1 *(npq1* + *Apxs*CVDE1). Values are average of replicate leaves and error bars represent standard error (n=6). ND, not detected. White bar represents illumination with 1000 µmol photons m^-2^ s^-1^ actinic light, black bar represents dark (with low measuring light) phase of the NPQ script.

## Discussion

### *Auxenochlorella* is a naturally occurring qE mutant

The slow kinetics of NPQ in WT *Auxenochlorella*, absence of an LHCSR-encoding gene in the genome, limited sequence similarity of the putative *Auxenochlorella* PSBSL1 to known PSBS, and the lack of NPQ phenotype in *psbsl1-1/psbsbl1-2* suggest that qE does not occur in this alga (Fig. 2a). While qE loss is unusual, it is not unprecedented; natural qE-deficient phenotypes have now been described in several green algae. The desert Trebouxiophyte green alga, *C. ohadii* lacks an *LHCSR* gene and encodes a *PSBS* gene that does not produce detectable levels of protein and correspondingly shows little to no qE (Levin et al., 2021) – similar to the *Auxenochlorella* phenotype observed here. Similarly, the siphonous green algae *Bryopsis* sp. and other Bryopsidales have independently lost LHCSR and PSBS and only exhibit slow NPQ kinetics (Christa et al., 2017; Mittila et al., 2026). Additionally, Bryopsidales can accumulate minor amounts of Zea, but they do not show the robust HL-induced VAZ-cycle conversion typically seen in organisms with qE or qZ in either HL- or LL-acclimated cells, consistent with Bryopsidales being natural qE and qZ mutants (Christa et al., 2017). Despite lacking both qE and a light-induced xanthophyll cycle, *Bryopsis* sp. exhibits slow reversible NPQ that is strikingly similar to that of *Auxenochlorella*.

Together, these examples show that loss of qE-related proteins has occurred at least three times within the green lineage. However, the ecological contexts of these losses are strikingly different: *C. ohadii* thrives in extreme desert soil crusts (Levin et al., 2021), *Bryopsis* sp. occupies rocky marine intertidal pools (Mattila et al., 2026), and *Auxenochlorella* was originally isolated from poplar tree sap (Krüger 1894; Craig et al., 2025). This suggests that qE loss is not correlated to a single environment or phylogenetic event. However, the losses do seem to recur in fast-growing, stress-tolerant algae, suggesting the loss of qE is tolerated or favored under specific selective pressures to prioritize traits such as rapid growth, heterotrophic growth, or broad stress resistance rather than maintaining fine-tuned qE.

Both *Bryopsis* sp. and *C. ohadii* seem to have evolved alternative photoprotective mechanism to tolerate high light. In *Bryopsis* sp., NPQ may arise from oxygen-dependent alternative electron pathways including flavodiiron-type activity, PTOX and mitochondrial respiration (Mattila et al., 2026). Unique alternative energy dissipation mechanisms have been observed in *C. ohadii,* such as dramatically reducing antenna size and chlorophyll *b*, and accumulation of lutein, Zea, Anthera, and β-carotene in HL (Levin et al., 2021). Given this precedent, our observation that *Auxenochlorella* lacks qE likely does not leave it undefended against HL but instead points to reliance on one or more alternative photoprotective routes such as pigment-based ROS scavenging, state transitions (qT), or unique cyclic electron flow and photosystem I control pathways that reduce excitation pressure on the photosynthetic apparatus.

### *Auxenochlorella* also does not exhibit robust qZ despite having a functional plant-type *Auxenochlorella* VDE1

Several lines of evidence support *Auxenochlorella* VDE1 as a plant-type de-epoxidase that may contribute to photoprotection in *Auxenochlorella* such as, the strong structural similarity to characterized plant-type VDE proteins (Fig. 3a, Fig. S3a), high Iso-seq transcript abundance during autotrophic growth (Craig et al., 2025), the DTT-sensitivity of NPQ under HL (Fig. 4a), the effect of VDE1 knockout on NPQ, and its ability to complement photoprotection when transiently expressed in *Nicotiana.* Although DTT caused a larger reduction in NPQ than deletion of *VDE1*, this difference may reflect the broader effects of DTT on multiple redox-dependent proteins involved in photosynthetic and photoprotective processes, including ascorbate peroxidase, glucose-6-phosphate dehydrogenase, and NADP-malate dehydrogenase (Neubauer and Yamamoto, 1994; Johnson, 1972; Yokochi et al., 2021). Although DTT treatment provides some evidence for the involvement of a DTT-sensitive de-epoxidation pathway, the *vde1* mutant allowed us to more specifically determine the contribution of *Auxenochlorella* VDE1 to NPQ and assess how *Auxenochlorella* compensates for the loss of a de-epoxidase. The modest reduction in NPQ observed in the *vde1* mutant indicates that *Auxenochlorella* VDE1 contributes only a minor fraction of total NPQ in *Auxenochlorella*, despite retaining genuine de-epoxidase activity, as demonstrated by its ability to rescue zeaxanthin accumulation and NPQ when expressed in *npq1 Nicotiana* (Fig. 8b and c). This suggests that *Auxenochlorella* relies primarily on VDE-independent mechanisms for photoprotection, a question we address in the following section.

*Auxenochlorella* does not exhibit the HL-induced conversion of Vio to Zea typically seen in model organisms like *Arabidopsis* and *Chlamydomonas* (Li et al., 2016). Vio and Zea levels remained relatively unchanged during HL exposure in WT *Auxenochlorella* (Fig. 4b and c; Fig. 6a and c; Fig. 7a and c). There are several possible explanations for these unusual VAZ pigment kinetics. One possibility relates to our Iso-Seq analysis, which showed that the vast majority of *VDE1* transcripts (96.5%) are truncated isoforms lacking the N-terminal domain, part of the lipocalin domain, and the transit peptide (Fig. S3a). This truncation may prevent both proper chloroplast localization and formation of an intact, catalytically active lipocalin domain, potentially resulting in a predominance of non-functional VDE1 protein in *Auxenochlorella*. This same truncation may also explain why *Auxenochlorella* VDE1 was able to complement the *Nicotiana npq1* mutant. The use of the *Arabidopsis* VDE1 transit peptide rather than the native *Auxenochlorella* sequence could have corrected the localization defect that affects the native, transit-peptide-lacking transcript. Additionally, the abundance of non-truncated *Auxenochlorella* VDE1 proteins may be insufficient to support substantial de-epoxidation of Vio and induction of qZ. Alternatively, physiological constraints of *Auxenochlorella*, such as differences in thylakoid pH dynamics in HL, may not support VDE activation (Hieber et al. 2012). Another possibility is that *Auxenochlorella* may be limited in ascorbic acid required for VDE activity (Müller-Moulé et al., 2002), for example due to limited transport of ascorbic acid into the thylakoid lumen or the ascorbic acid pool being depleted faster by the competing ascorbate peroxidase (Neubauer et al.,1994). Further investigation of the VAZ cycle in *Auxenochlorella* will be the subject of future studies.

Although VDE1 does not appear to mediate canonical qZ, deletion of *VDE1* resulted in altered carotenoid composition and photoprotective responses. Differences observed in *vde1* and *vde1 cvde1* were increased Anthera and lutein. While the removal of VDE1 abolished detectable levels of Anthera and Zea in *Nicotiana* and *Arabidopsis npq1* mutants (Fig. 8b) (Li et al., 2016; Lee et al., 2024), both *vde1* and *vde1 cvde1* exhibited increased Anthera (Fig. 7b). The accumulation of Anthera in *vde1* may arise from VDE-independent Zea and Anthera biosynthesis pathways; Zea can be formed through hydroxylation of β-carotene and Anthera from Zea through the activity of ZEP (Lohr et al. 2005). The opposing effects on Anthera accumulation by DTT (decreased Anthera) and that by loss of VDE1 and CVDE1 in the mutants *vde1* and *vde1 cvde1* (increased Anthera) could be explained by DTT negatively affecting the VDE-independent Anthera (and Zea) forming pathway. However, it has been shown that ZEP is not directly downregulated through disulfide reduction by DTT but is sensitive to H_2_O_2_ (Netto and Stadtman, 1996; Pérez and Maureira, 2003; Holzmann et al., 2022). H_2_O_2_ is a byproduct of the autoxidation reaction of DTT in aerobic conditions, although how much H_2_O_2_ is produced under our experimental conditions is not known.

The *vde1 cvde1* mutant exhibited decreased Vio and increased lutein suggesting a redistribution of flux upstream in carotenoid metabolism toward the lutein-producing branch (Fig. 6a and f). This change in carotenoid allocation may indicate that loss of VDE1 affects carotenoid allocation independently of detectable cycling activity in *Auxenochlorella*. The basis for this shift remains unclear and may reflect a role for VDE1 beyond xanthophyll cycling, such as a constitutive effect on carotenoid pool partitioning that is not captured by measuring cycle induction alone. Taken together, the altered carotenoid profiles of the *vde1* and *vde1 cvde1* mutants demonstrate that VDE1 contributes to carotenoid homeostasis and photoprotective responses in *Auxenochlorella*. However, the residual xanthophyll pigments and the absence of a canonical VDE-dependent qZ response in WT, suggests that VDE1 may have a photoprotective role independent of qZ mediation in this system.

The small contribution of VDE1 in alternative photoprotection is further supported by the PAM and TCSPC phenotypes observed in the *vde1* mutants. The *vde1* mutant had approximately a 25% reduction in fluorescence yield when imaged via PAM (Fig. 5a). However, the excited state decay kinetics revealed that the modest quenching deficit between the *vde1* mutant and the reference strain was much smaller than what was observed via PAM (Fig. 5a and c). NPQτ increased over multiple cycles of light and dark in all genotypes, but the *vde1 cvde1* deviated less from the reference in the second cycle compared to the first, and this is consistent with elevated accumulation of photodamage or slow photoprotective processes during HL exposure in the mutant. The difference in the magnitude of the NPQ observed by PAM and TCSPC suggests that the reduced fluorescence yield measured by PAM is not solely attributable to a decrease in the excited state chlorophyll lifetimes, but may also reflect changes in baseline F_m_, closed reaction-center populations, or genotype-dependent differences in pigment– antenna organization. Together, these data support a model in which the loss of *VDE1* and *CVDE1* results in sensitivity to repeated HL exposure, which we speculate is associated with changes in carotenoid composition. Because these mutations did not abolish the accumulation of de-epoxidized xanthophylls (Zea and Anthera) or reversible NPQ, *Auxenochlorella* NPQ must be comprised of one or multiple alternative photoprotective pathways and not qZ.

### The cryptic role of *Auxenochlorella* CVDE1 *in vivo*

Like *Auxenochlorella* VDE1, the sequence similarity to algal CVDEs and complementation of *Nicotiana npq1* mutant led us to hypothesize the involvement of *Auxenochlorella* CVDE1 in qZ NPQ.

However, deletion of *CVDE1* resulted in only minor changes in NPQ, NPQτ (Fig. 5), and VAZ pigment abundance or composition (Fig. 6a-c). Much of the HL-related phenotype in a *cvde1* background was observed only when combined with *vde1*, suggesting that *Auxenochlorella* CVDE1 makes a minor contribution *in vivo* and that *Auxenochlorella* VDE1 activity may be compensating in the *cvde1* mutant. Overall, *Auxenochlorella* CVDE1 does not contribute to changes in VAZ pigment composition or NPQ, despite the observation that *Auxenochlorella CVDE1* encodes a functional de-epoxidase.

We did not investigate whether *Auxenochlorella* CVDE1 has a CruP-like function *in vivo* in *Auxenochlorella.* The precise function of CruP remains unresolved, although previous studies have associated CruP with the level of β-carotene-5,6-epoxide and protection against ROS accumulation (Bradbury et al., 2012). In *A. thaliana*, loss of CruP results in increased β-carotene-5,6-epoxide, decreased β-carotene, and elevated ROS levels (Bradbury et al., 2012), pointing to a possibility that CruP is a β-carotene-5,6-epoxide de-epoxidase as suggested by Li et al. (2016). In contrast, the *cvde1* mutant accumulated higher β-carotene while maintaining fluorescence yield and NPQτ similarly to WT, and there was no increase in slow, photoinhibitory type NPQ or NPQτ (Fig. 5b and d). Additionally, we did not find an alternative CruP-like homolog in *Auxenochlorella*, suggesting that this organism may lack the CruP protein.

### Conclusion

Our study demonstrates that *Auxenochlorella* is a natural qE-deficient alga lacking LHCSR and a *bona fide* PSBS. Despite encoding genes for functional VDE and CVDE our results showed that *Auxenochlorella* is also naturally qZ-deficient. The deletion of one or both de-epoxidases did not eliminate the bulk of reversible NPQ induction and relaxation in *Auxenochlorella*. Most importantly, the inverse relationship between Vio and Zea typically observed during NPQ induction and relaxation, reflecting VDE- and ZEP-mediated xanthophyll-cycle activity, was absent in WT *Auxenochlorella*. The kinetics of NPQ relaxation would seem to exclude photoinhibition (qI) as the only NPQ component, although the minor increase in NPQ with slow kinetics over time suggests that qI does occur in HL (Fig. 4).

The exact photoprotective mechanism in *Auxenochlorella* still remains unidentified but may comprise a combination of qT, qH, cyclic electron flow and photosystem I control pathways, or ROS scavenging pathways. The qH component of NPQ seems unlikely, as the *Auxenchlorella* genome does not encode Lipocalin in the Plastid (LNCP), Suppressor of Quenching 1 (SOQ1), or Relaxation of QH 1 (ROQH1). On the other hand, the involvement of qT can be tested by generating mutants in the putative gene encoding STT7. This study highlights the importance of considering genetic and lineage diversity when characterizing NPQ pathways in photosynthetic organisms. *Auxenochlorella* provides an interesting system to study NPQ in an organism lacking the canonical qE and qZ mechanisms, suggesting other NPQ pathways may play important roles across the tree of life.

## Acknowledgements

We thank Elina Patel for assisting with TCSPC data collection.

## Author contributions

K. C., J. H., L. G., R. I., C. B., and S. W. performed research; J. K. and J. M. designed and generated the *psbs*l1-1 and *psbsl1-1/psbsl1-2* lines; D. C. designed and generated the THIC_ref_ line; K. C., S. W., J. H., L. G, K. K. N., and G. R. F. designed research; K. C., J. H., L. G., S. W., C. B., and K. K. N. analyzed data and wrote the paper; all authors discussed the results and commented on the manuscript.

## Data Availability Statement

The data supporting the findings of this study are available within the article and its supplementary materials. Plasmids used for transient expression in *Nicotiana* and plasmids for generating *Auxenochlorella* mutants can be found on Addgene.

