## Supplementary Figure 1 for "A Natural qE- and qZ-Deficient Alga Retains Functional Plant-Like and Chlorophycean Violaxanthin De-Epoxidases"

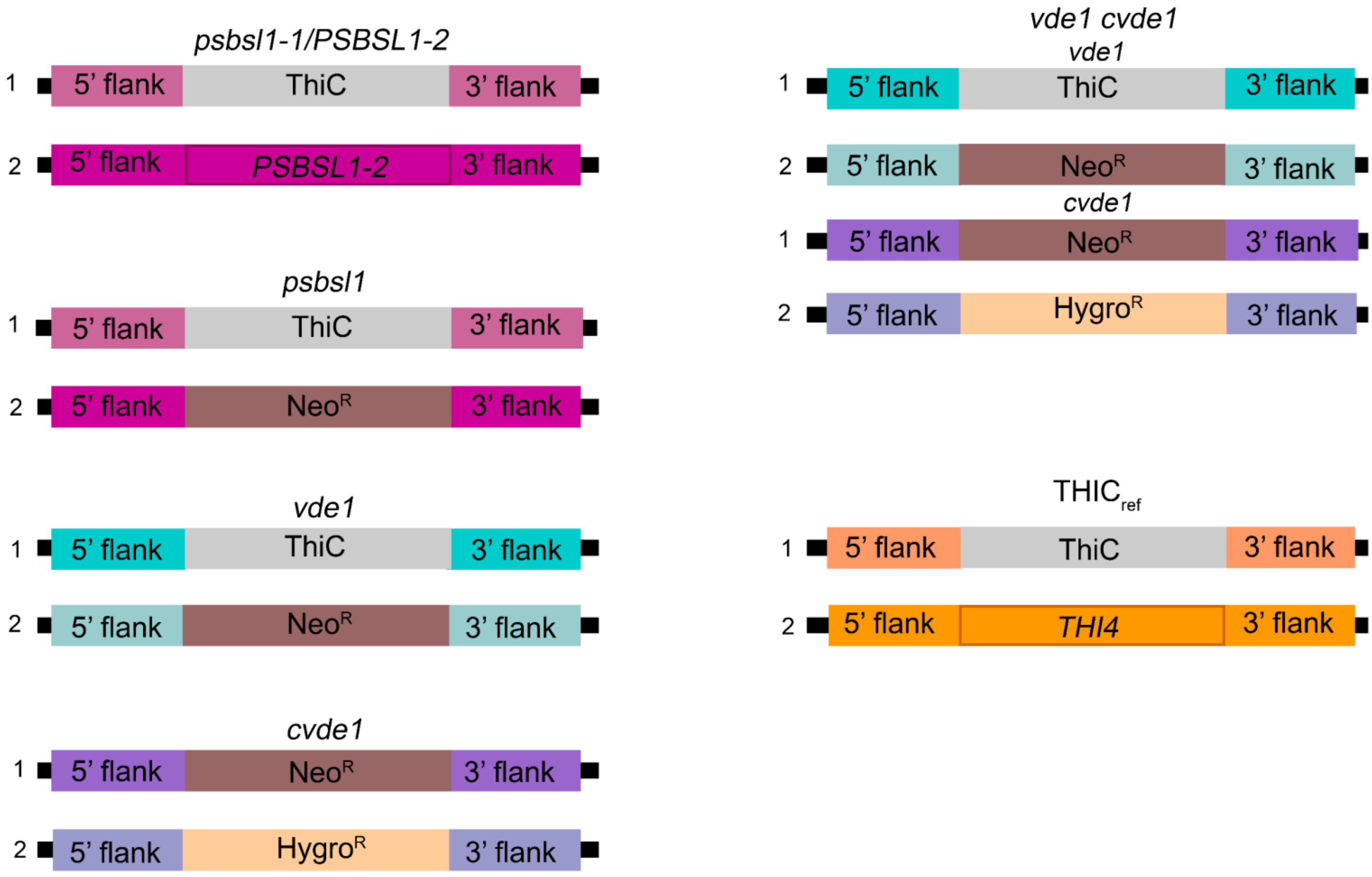


**Supplementary Figure 1. Schematic summary of the target loci in the mutants generated in this study.** Each allele was targeted with a construct containing a selection marker flanked by homologous arms on the 5' and 3' side. Fragments not to scale.
