## Supplementary Figure 2 for "A Natural qE- and qZ-Deficient Alga Retains Functional Plant-Like and Chlorophycean Violaxanthin De-Epoxidases"

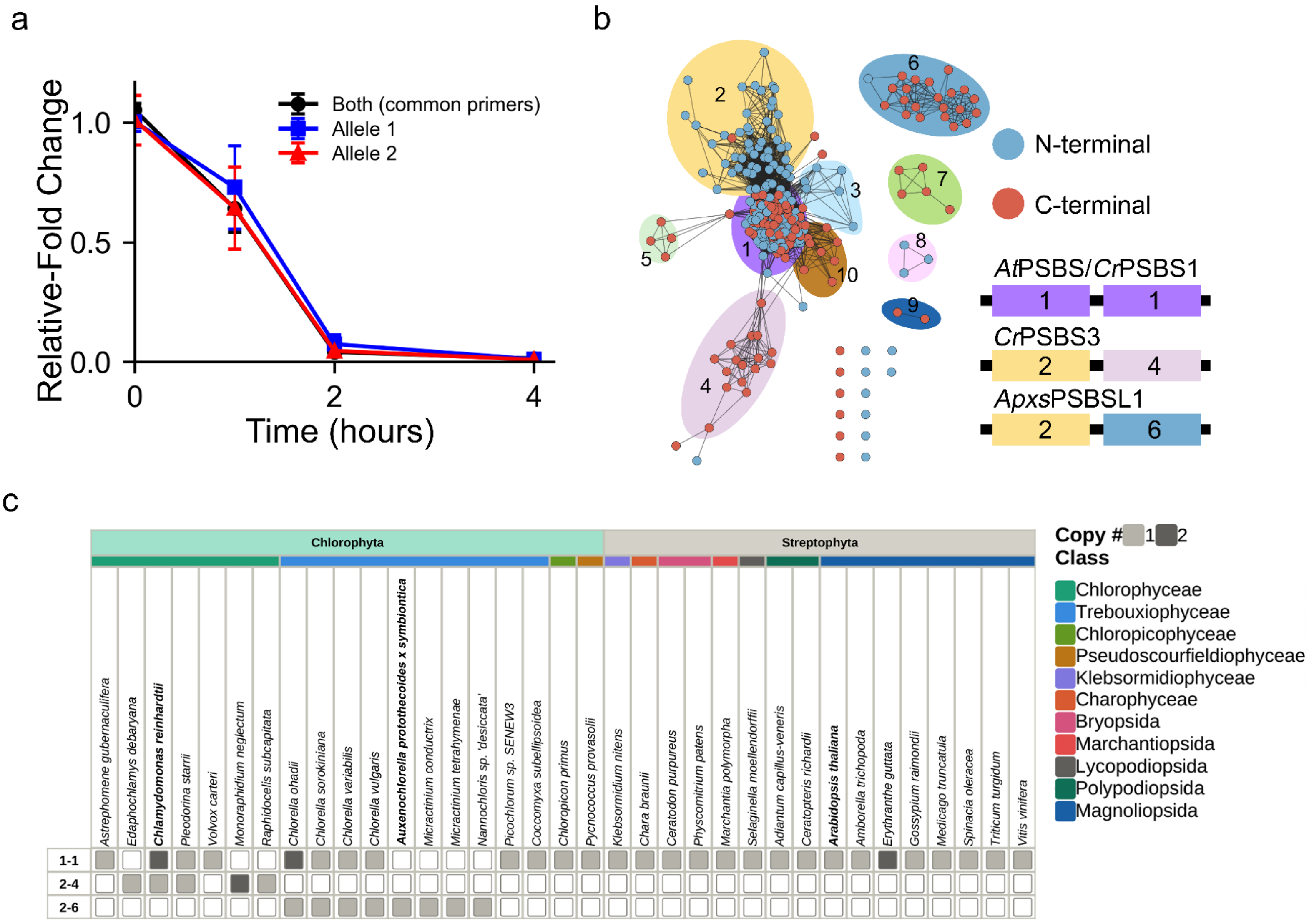


**Supplemental Figure 2. *Auxenochlorella* PSBSL1 has divergent pairs of N- and C-terminal helices. a)** *Auxenochlorella PSBSL1* transcript fold change when transferred from LL to HL. Values represent averages of biological replicates and standard error shown as error bars (n=3). **b)** SSN analysis organized according to identified clusters. Nodes are colored by N- or C-terminal region according to color key, and background shades and numbering refers to identified clusters of helices pairs. Cartoon diagrams correspond to the N and C-terminal cluster combinations of *Arabidopsis* PSBS (*At*PSBS), *Auxenochlorella* PSBSL1 (*Apxs*PSBSL1), *Chlamydomonas* PSBS1 (*Cr*PSBS1*)* and PSBS3 (*Cr*PSBS3) homologs identified in the SSN. **c)** Distribution of identified N- and C-terminal cluster combinations in plants and green algae from panel a. Cluster combinations are written as N-terminal cluster number followed by C-terminal cluster number.
