## Supplementary Figure 3 for "A Natural qE- and qZ-Deficient Alga Retains Functional Plant-Like and Chlorophycean Violaxanthin De-Epoxidases"

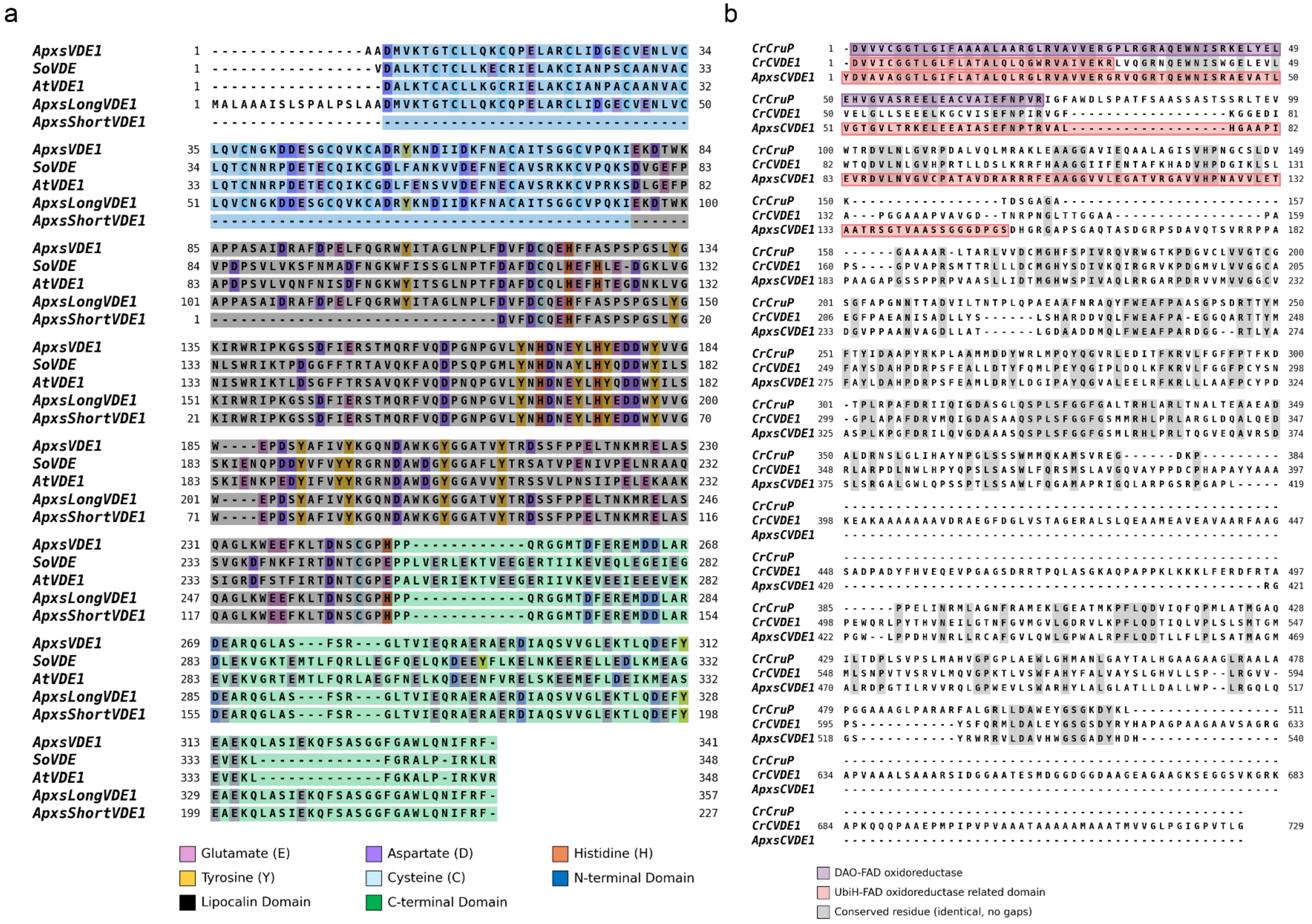


**Supplementary Figure 3. Protein alignment and domain comparison in *Apxs*VDE1 and *Apxs*CVDE1.** Protein alignments for VDE were generated from *Spinacia oleracea* (UniProt: Q9SM43), *Arabidopsis* VDE1 (AT1G08550), *Auxenochlorella* VDE1 (UTEX250_B10.26655), and the long and truncated Iso-seq reads from Craig et al., 2025 dataset (SRA: SRR32007620). Protein alignments for CVDE were generated from *Auxenochlorella* CVDE1 (UTEX250_B11.34560) and *Chlamydomonas* CVDE1 (Cre04.g221550.t1.2) and CruP (Cre06.g264250). Transit peptides were predicted using TargetP 2.0 (Almagro Armenteros et al., 2019) and mature proteins were generated from removing the transit peptide. Sequence alignments were generated using the EMBL-EBI MUSCLE alignment tool with default parameters (Edgar, 2004; Madeira et al., 2024). Conserved domains were identified using the NCBI Conserved Domain Database search function (Wang et al., 2023 ). Jalview v2.11.5.2 (Waterhouse et al., 2009) was used to visualize the resulting alignment. **a)** Annotated alignment of the mature *Arabidopsis* VDE1 *(At*VDE1)*, S. oleracea* VDE *(So*VDE)*, Auxenochlorella* VDE1 (*Apxs*VDE1), long (*Apxs*LongVDE1) and truncated Iso-seq reads (*Apxs*ShortVDE1) . Domain and amino acid annotations are based on residues and domains of known significance in *So*VDE. **b)** Annotated alignment of the mature CVDE1 from *Auxenochlorella (Apxs*CVDE1), *Chlamydomonas (Cr*CVDE1), and *Chlamydomonas CruP(Cr*CruP). Domain is based on NCBI domain annotations.
