## Supplementary Figure 4 for "A Natural qE- and qZ-Deficient Alga Retains Functional Plant-Like and Chlorophycean Violaxanthin De-Epoxidases"

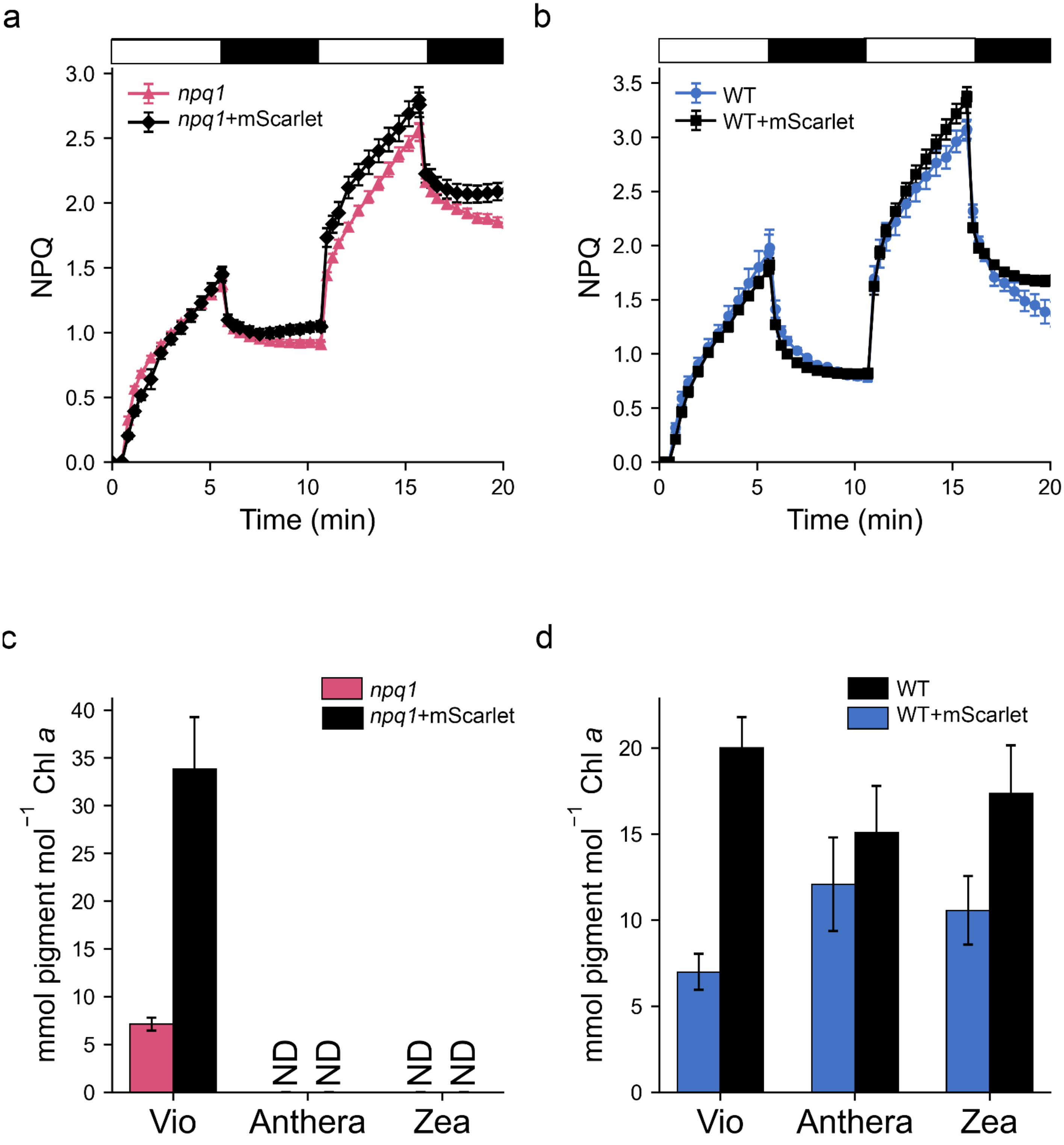


**Supplementary Figure 4. mScarlet does not affect NPQ in the *npq1* or WT *Nicotiana.*** NPQ of **a)** *npq1* versus *npq1* expressing mScarlet (*npq1* + mScarlet) and **b)** WT versus WT expressing mScarlet (WT + mScarlet). VAZ pigments in **c)** *npq1* versus *npq1* expressing mScarlet and **d)** WT versus WT expressing mScarlet. Pigments were extracted after 30 minutes of 1000 µmol photons m$^{-2}$ s$^{-1}$ actinic light. Values are average of replicate leaves and error bars represent standard error (n=6). ND, not detected.
